# Spatially resolved host–fungal transcriptomics profiling by fungal-DBiT

**DOI:** 10.64898/2026.10.19.760099

**Authors:** Xiaohui Yu, Fu Gao, Deguang Song, Gaocai Li, Dejiang Wang, Shenjie Wang, Xiangjun Di, Shujie He, Liyun Lin, Junyi Xu, Di Zhang, Yu Jin, Jing Du, Noah W. Palm, Yang Liu

## Abstract

Hosts and their associated fungi coexist within a shared tissue environment, engaging in dynamic interactions that influence immune regulation, tissue homeostasis, and disease progression. Dissecting these interactions requires spatially resolved and unbiased approaches capable of capturing gene expression from both host and fungal cells within intact tissue architecture. Here, we present fungal-DBiT, a spatial transcriptomic platform that enables simultaneous, high-resolution mapping of host and fungal RNAs, directly in tissue sections. Using fungal-DBiT, we profiled the spatial distribution of host and fungal transcripts in a fungal infection mouse model, and review pronounced spatial heterogeneity in fungal localization and activity, accompanied by region-specific host transcriptional responses. We further applied fungal-DBiT to intestinal tissues under physiological conditions, demonstrating its sensitivity in detecting low-abundance fungal species in the absence of overt inflammation. Together, these results demonstrate the ability of fungal-DBiT to resolve host–fungal interactions in situ and highlight its utility for studying mycobiome-associated immune responses and disease processes with spatial precision.

## Background

Invasive fungal infections represent a major and growing global health burden, causing more than three million deaths annually worldwide^1, 2^. Although fungi comprise a minor fraction of the human microbiome, they exert outsized effects on host immunity and disease progression, particularly in immunocompromised populations, and their ecological range is expanding with climate change^3, 4^. Despite their clinical importance, fungi remain poorly characterized within mammalian tissues, in part due to a lack of technologies capable of resolving fungal biology in situ.^5, 6^ Most microbiome and transcriptomic approaches profile fungal communities in bulk, obscuring spatial organization and local host responses, while existing imaging-based methods provide limited molecular depth.^7, 8^

Spatial transcriptomics has emerged as a transformative approach for interrogating tissue organization and host–pathogen interactions by coupling gene expression profiling with spatial context.^9–11^ However, current spatial transcriptomic technologies are largely optimized for mammalian host cells and have not been systematically adapted for fungal organisms. Technical challenges, including rigid fungal cell walls, inefficient RNA capture, have limited the application of spatial transcriptomics to fungi in mammalian systems.^12, 13^ As a result, spatial fungal microstructures, fungal burden heterogeneity, and localized host immune architectures during infection remain largely uncharacterized. Existing fungal spatial studies have focused primarily on plant systems or rely on commercial platforms with limited flexibility and throughput^14, 15^.

Here, we present fungal-DBiT (deterministic barcoding in tissue for fungal analysis), a spatial transcriptomics platform specifically optimized for fungal biology that enables simultaneous, spatially resolved profiling of fungal and host transcriptomes in intact mammalian tissues. Fungal-DBiT integrates targeted fungal cell wall permeabilization with orthogonal microfluidic barcoding to achieve efficient fungal RNA capture while preserving tissue architecture. We demonstrate robust co-detection of fungal and host transcripts across multiple mouse organs, validate fungal localization using 18S rRNA fluorescence in situ hybridization, and reveal organ-specific host immune responses alongside distinct fungal spatial patterns. By extending spatial transcriptomics to fungi in mammalian tissues, fungal-DBiT provides a generalizable framework for studying host–fungal interactions, polymicrobial infections, and fungal pathogenesis in situ.

## Results

### The overall design of fungal-DBIT

To realize spatially resolved analysis of fungal and host transcriptomes within intact tissues, we developed fungal-DBiT, a spatial transcriptomics platform specifically optimized for fungal biology. Fungal-DBiT adapts deterministic barcoding in tissue for spatial omics sequencing (DBiT-seq)^16^ by incorporating fungal cell wall removal and optimized permeabilization to allow efficient in situ capture of fungal RNAs alongside host transcripts (**Figure 1**). Following tissue fixation, reverse transcription is performed directly on tissue sections, after which two rounds of orthogonal microfluidic barcoding introduce spatial barcode A and barcode B through sequential ligation, thereby encoding two-dimensional spatial coordinates at the transcript level together with unique molecular identifiers. Barcoded cDNAs are subsequently released, amplified, and prepared for sequencing (**Figure 1a**). A dedicated computational pipeline performs quality control, dual alignment to host and fungal reference genomes, reads classification, and spatial count matrix generation, enabling integrated analysis of fungal load, spatial distribution, and host transcriptional responses (**Figure 1b**).

**Figure 1.**
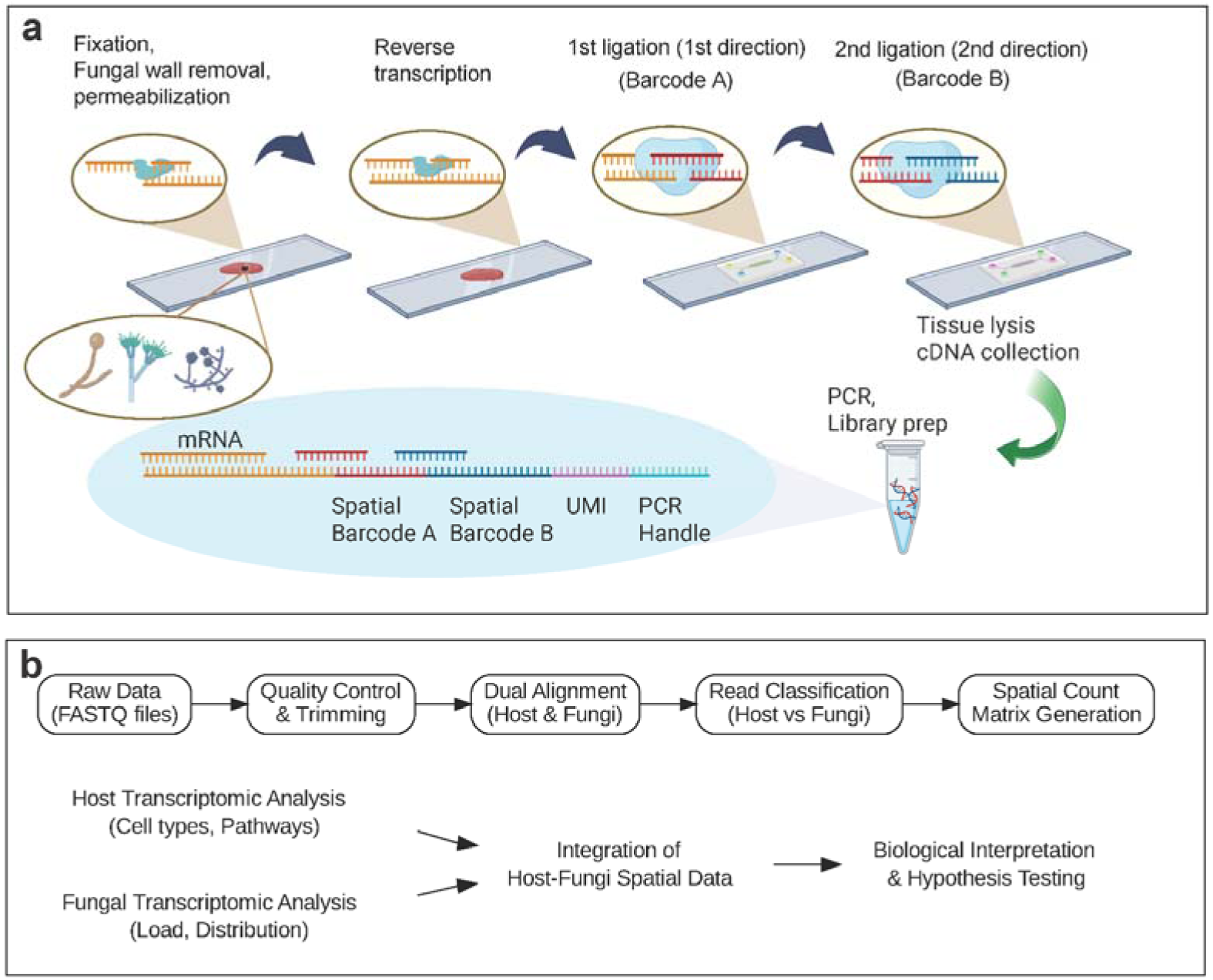
Experimental and computational workflow of fungal-DBiT. a, Schematic of the fungal-DBiT experimental workflow for spatially resolved host–fungal transcriptomics. Tissue sections containing fungal cells are first fixed, followed by fungal cell wall removal and tissue permeabilization to enable RNA access. Reverse transcription is performed in situ, after which two sequential orthogonal microfluidic barcoding steps introduce spatial barcode A and spatial barcode B through ligation in perpendicular directions. Barcoded cDNAs containing spatial barcodes, UMI region, and PCR handles are released by tissue lysis, followed by PCR amplification and library preparation for sequencing. b, Overview of the fungal-DBiT computational analysis pipeline. Raw sequencing reads undergo quality control and trimming, followed by dual alignment to host and fungal reference genomes. Reads are classified as host or fungal and spatially resolved count matrices are generated. Downstream analyses include host transcriptomic profiling (cell type annotation and pathway analysis), fungal transcriptomic analysis (fungal load and spatial distribution), and integrated host– fungal spatial analysis for biological interpretation and hypothesis testing.

### Fungal-DBiT profiling of heart tissue from a proof-of-concept mouse model

To evaluate the performance of Fungal-DBiT in complex mammalian tissues, we established an experimental mouse model using a defined mixture of seven fungal species (**Figure 2a**, see Methods), including members of the *Candida* and *Aspergillus* genera (**Table S1**). The mixture included both yeasts and filamentous fungi, representing distinct fungal morphologies. This experimental system was designed as a technical proof-of-concept, analogous to the use of defined cell mixtures for benchmarking single-cell sequencing technologies, to assess the feasibility, sensitivity, and species-resolving capability of Fungal-DBiT in the context of host tissues. Importantly, the model was not intended to recapitulate naturally occurring fungal infections or their pathophysiological processes, but rather to provide a controlled setting for evaluating simultaneous spatial profiling of host and multiple fungal transcriptomes.

**Figure 2.**
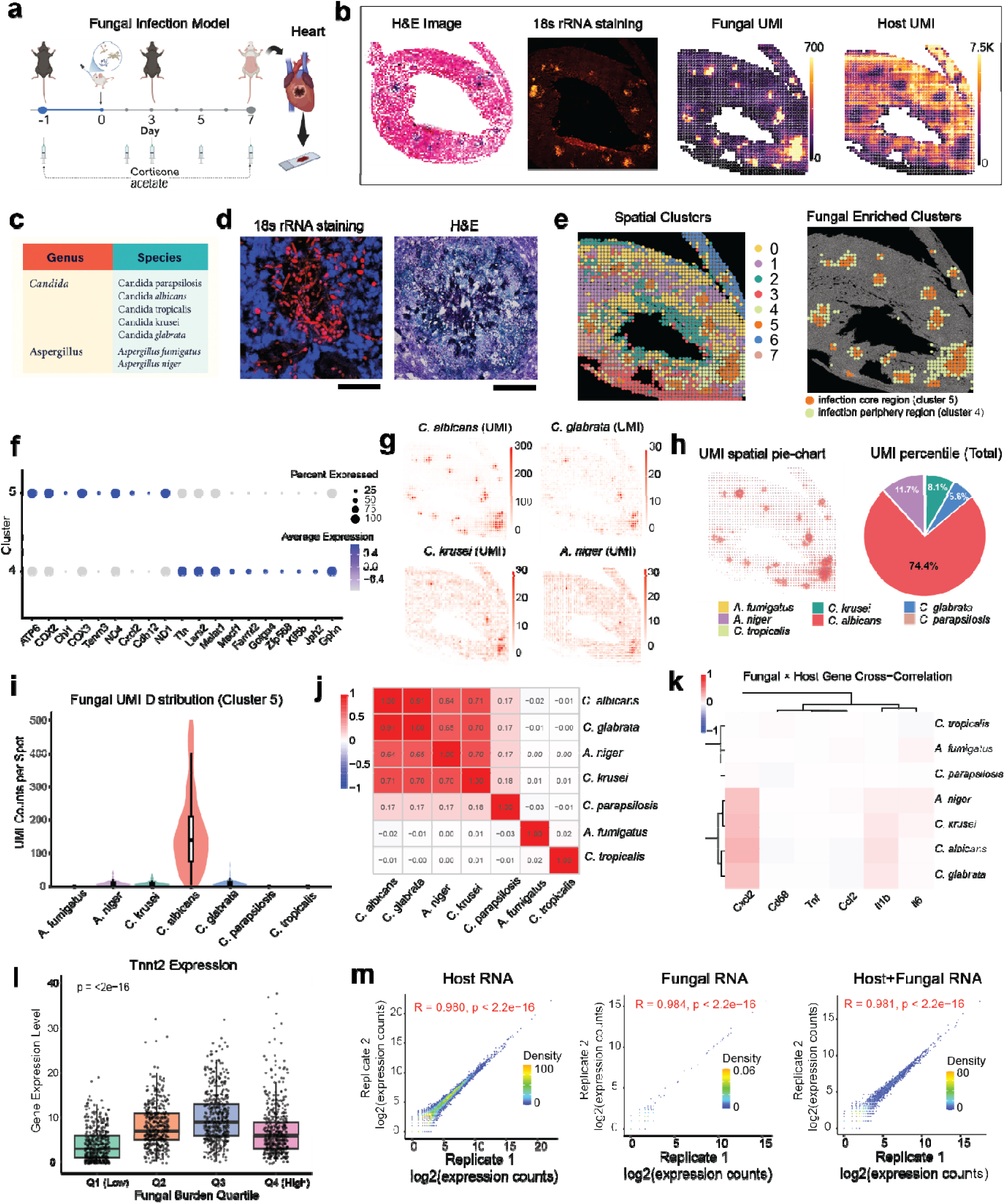
Spatially resolved host–fungal transcriptomics profiling of infected mouse heart using fungal-DBiT. a, Schematic of the in vivo fungal infection model and experimental workflow. Mice were treated with cortisone acetate, followed by systemic fungal infection. Heart tissues were harvested at indicated time points and processed for fungal-DBiT spatial transcriptomic profiling. b, Spatial maps from a heart section, including H&E staining, fungal 18S rRNA staining, fungal UMI counts, and host UMI counts. c, Taxonomic coverage of fungal-DBiT, showing detected fungal genera and species included in the analysis. d, High-resolution fluorescence imaging of fungal 18S rRNA staining (left) and corresponding H&E staining (right). e, Unsupervised spatial clustering of fungal-DBiT data identifies distinct tissue clusters, with highlighted fungal-enriched clusters corresponding to infection core and peripheral regions. f, Dot plot showing differential host gene expression across spatial clusters. g, Spatial expression maps of representative fungal species (UMI counts) across the tissue section. h, Spatial pie-chart visualization of fungal species composition across tissue spots (left) and overall fungal UMI composition across the section (right). i, Violin plot showing distribution of fungal UMI counts per spot for each fungal species within the cluster 5. j, Pairwise correlation matrix of fungal species UMI abundance across spatial spots, revealing co-occurrence and exclusivity patterns among fungal taxa. k, Heatmap of cross-correlations between fungal species abundance and host immune gene expression, highlighting species-specific host response signatures. l, Host gene (Tnnt2) expression stratified by fungal burden quartiles, demonstrating association between fungal load and host transcriptional changes (Wilcoxon test). m, Reproducibility of fungal-DBiT across biological replicates, shown by correlation of log-transformed expression counts for host RNA, fungal RNA, and combined host–fungal RNA profiles.

Mice were maintained under immunosuppressed conditions for one week before tissue collection (**Figure 2a**). Heart and kidney tissues were subsequently collected for spatial profiling.

We first performed fungal-DBiT using 50 μm resolution chips (spot size: 50×50 μm) and on average obtained 5000-6000 UMIs per spot (host and fungal together), covering a total of 20000-30000 genes across all the spots. We detected 0-1000 UMIs and 0-200 genes for fungal RNAs (**Figure 2b, Figure S1a**) in each spatial spot, depending on the fungal abundance in each spot. We also conducted general 18S rRNA staining using a neighboring tissue section and showing similar patterns that match with fungal UMI distribution (**Figure 2b**). Interestingly, we observed an inverse relationship between fungal and host UMI distributions, such that regions with higher fungal abundance exhibited markedly reduced host UMI counts. This pattern is consistent with fungal-induced host cell loss, potentially through apoptosis or tissue damage.^17^ To independently validate this observation in higher resolution, we compared 18S rRNA fluorescence in situ hybridization with corresponding H&E-stained tissue sections(**Figure 2d**). Consistent with the spatial transcriptomic data, regions with high fungal signal displayed reduced host cell density, supporting a spatially localized depletion of host cells in areas of fungal enrichment.

We next performed unsupervised clustering using combined host and fungal gene expression profiles from the heart tissue and identified a total of eight spatial clusters (**Figure 2e**). Based on established host marker genes, we assigned cell-type identities to the host cell populations across clusters (**Figure S1b–d**). Among these clusters, two exhibited marked enrichment of fungal UMIs, with cluster 4 spatially surrounding cluster 5 across most regions of the tissue. We therefore designated cluster 4 as the “infection periphery region” and cluster 5 as the “infection core region”. Notably, cluster 5 was also enriched for myeloid cell markers, suggesting an early host immune response associated with fungal invasion. To further characterize host responses associated with fungal localization, we compared host gene expression between clusters 4 and 5 (**Figure 2f and S2**). Cluster 4 showed reduced expression of mitochondrial and contractile genes, including mt-*Atp6*, *Cox1*, *Cox3*, *mt-Cytb*, and *mt-Nd1*, whereas cluster 5 retained relatively higher expression of these markers. In contrast, genes involved in nuclear regulation and transcriptional control, such as *Cmss1*, *Cdk8*, and *Malat1*, were enriched in cluster 4. These patterns indicate spatially localized host transcriptional differences associated with fungal presence. We additionally identified host genes whose expression correlated with the spatial distribution of fungal signals (**Figure S2b**). Nuclear regulatory genes (*Cmss1*, *Cdk8*, *Malat1*) were consistently enriched in fungal-dense regions, whereas mitochondrial and cardiomyocyte structural genes (*Lars2*, *Cox1*, *mt-Cytb*, *Ttn*) were reduced, supporting spatially localized host transcriptional reprogramming in response to fungal colonization.

We next mapped the spatial distribution of seven fungal species across infected heart tissue (**Figure 2g**, **2h and S3**). Among the detected species, *C. albicans* exhibited the highest UMI abundance and the most pronounced spatial enrichment, forming discrete high-density regions. Other species, including *C. glabrata*, *C. krusei*, and *A. niger*, were detected but at lower abundance. These patterns indicate that fungal colonization within cardiac tissue is spatially restricted and varies substantially across species. Quantitative analysis of fungal composition across all spatial spots further demonstrated the dominance of *C. albicans*, which accounted for the majority of fungal UMIs (74.4%), whereas other species contributed smaller fractions of the total fungal transcriptome (**Fig. 2h**). The representative genes for each species were shown in **figure S4**. Cluster 4 and 5 show similar fungi composition, but cluster 5 have much higher overall fungi load (**Figure 2i, S3b and S3c**). We also performed bulk RNA-seq on adjacent heart tissue sections and observed strong concordance with our spatial RNA-seq data (r = 0.847, **Figure S3d and 3e**). Pairwise correlation analysis of fungal species revealed distinct patterns of spatial co-occurrence across the infected heart tissue (**Figure 2j**). *Candida* species, including *C. albicans*, *C. glabrata*, and *C. krusei*, showed strong positive correlations with each other, indicating shared or overlapping spatial niches. Cross-correlation analysis between fungal abundance and host gene expression further revealed species-specific associations with host immune-related genes (**Figure 2k**). In particular, *Candida* species showed modest positive correlations with chemokine and inflammatory markers such as *Cxcl2* and *Il1b*, whereas other fungal taxa displayed minimal or no association with these host genes.

Stratification of spatial spots by fungal burden revealed a significant association between local fungal abundance and host gene expression (**Figure 2l and Fig.S5**). Expression of *Tnnt2*, a cardiomyocyte contractile marker, varied across fungal burden quartiles, with reduced expression observed in spots with higher fungal load. This pattern is consistently observed across additional contractile and cytoskeletal genes (*Myh6, Myl3, Tnni3, Acta2, Des*), indicating a coordinated activation of muscle-like or stress-responsive programs in higher-Q regions. In parallel, genes involved in calcium handling and metabolic activity (*Ryr2, Atp2a2, Camk1d*) also show increasing expression toward Q3/Q4, suggesting enhanced energetic and signaling demands in these spatial niches. This relationship suggests that increased local fungal presence is associated with altered host gene expression in a burden-dependent manner, highlighting the ability of Fungal-DBiT to link spatial microbial signals with host transcriptional changes.

Reproducibility of Fungal-DBiT was assessed by comparing gene expression measurements between biological replicates for host RNA, fungal RNA, and the combined host–fungal transcriptome (**Figure 2m**). Strong correlations were observed across replicates for host transcripts (Pearson’s R = 0.980), fungal transcripts (R = 0.984), and combined datasets (R = 0.981), indicating high consistency despite differences in RNA abundance and composition.

### Spatial profiling of infected heart tissue in high resolution

We next performed spatial profiling of infected heart tissue using a 25 µm-resolution chip (**Figure 3**). Consistent with results obtained using the 50 µm chip, we observed pronounced spatial heterogeneity in fungal colonization, with fungal RNAs detected in discrete tissue regions rather than uniformly distributed across the myocardium (**Figure 3a and 3b**). Unsupervised clustering identified the same major host cell types as those resolved at 50 µm resolution (**Figure 3c, Figure S6a and S6b**), and clearly delineated fungal infection core and periphery regions. Comparative analysis between these regions revealed gene-expression trends consistent with those observed at lower resolution (**Figure 3d**).

**Figure 3.**
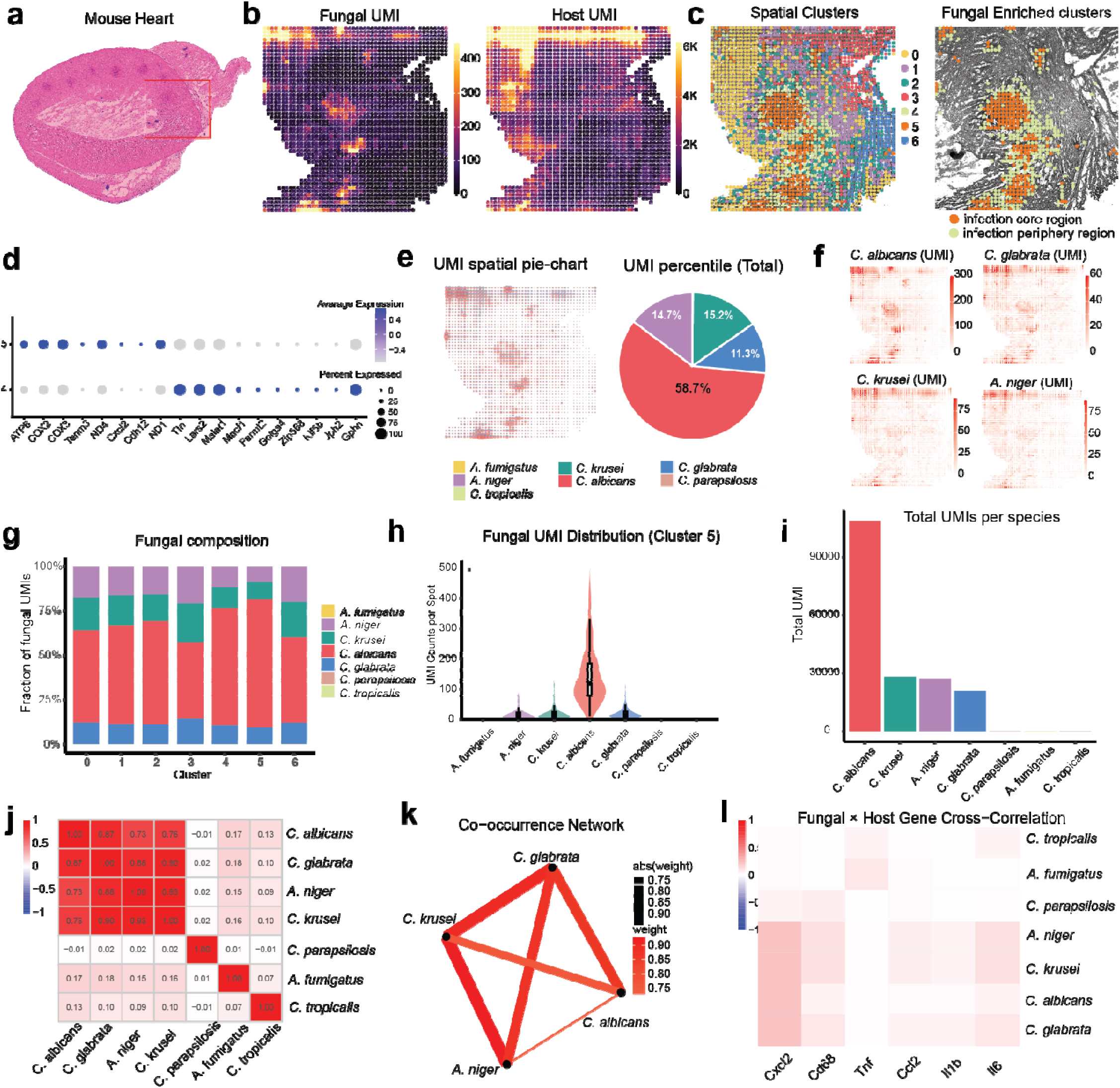
Spatial profiling of fungal infected heart tissue with higher resolution. a, H&E staini g of an infected heart section, with the region profiled by fungal-DBiT indicated. b, Spatial maps of fungal UMI counts and host UMI counts across the tissue section. c, Unsupervised spatial clustering of fungal-DBiT data identifies distinct tissue niches, including clusters enriched for fungal transcripts. d, Dot plot showing differential host gene expression across spatial clusters 4 and 5. e, Spatial pie-chart representation of fungal species composition across tissue spots. f, Spatial expression maps of representative fungal species (UMI counts), illustrating species-specific localization patterns within infected tissue. g, Overall fungal species composition summarized as the fraction of total fungal UMIs per cluster. h, Violin plot showing the distribution of fungal UMI counts per spot for individual fungal species within a representative infection-enriched cluster. i, Total fungal UMI counts per species across the tissue section. j, Pairwise correlation matrix of fungal species abundance across spatial spots, revealing co-occurrence patterns among fungal taxa. k, Network visualization of fungal species co-occurrence relationships inferred from spatial correlations. l, Heatmap of cross-correlations between fungal species abundance and host immune-related gene expression, highlighting species specific host response signatures.

Fungal-enriched clusters corresponding to infection core regions exhibited pronounced spatial heterogeneity in fungal burden, with total fungal UMI counts varying substantially across spots and reaching several hundred UMIs per spot in localized hotspots (**Figure 3e**). Species-resolved analysis revealed that *C. albicans* are the predominant fungal species within these regions, accounting for approximately 58.7% of total fungal UMIs and reaching local maxima of ∼300 UMIs per spot, consistent with focal overgrowth. In contrast, other *Candida* species, including *C. tropicalis* (∼15.2%), *C. glabrata* (∼11.3%), and *C. krusei* (∼14.7%), are detected at lower overall abundance and exhibited more spatially diffuse and less sharply defined distributions (**Figure 3f–h**). Pairwise correlation analysis further demonstrated strong positive spatial correlations among *Candida* species (Pearson’s r ≈ 0.7–0.9), suggesting partial co-localization or shared infection niches, whereas correlations involving *Aspergillus* species were weak (r ≈ 0-0.2), indicating distinct or non-overlapping spatial patterns (**Figure 3i,j**).

Integrative analysis of fungal and host transcriptomes further revealed localized associations between fungal-enriched regions and host inflammatory gene expression, enabling spatially resolved interrogation of host– fungal interactions (**Figure 3k,l**). Together, these results demonstrate that Fungal-DBiT provides reproducible, species-resolved spatial quantification of fungal communities and their associated host transcriptional responses in intact cardiac tissue at higher spatial resolution.

### Spatial profiles of host and fungal transcriptomes in infected kidneys

Kidney tissue from the infected model was similarly profiled using Fungal-DBiT, revealing pronounced spatial heterogeneity in fungal colonization, with fungal RNA detected in discrete regions rather than uniformly across the tissue (**Figure 4a**). Compared with heart tissue, fungal infection in the kidney was more broadly distributed. Across the tissue, we detected an average of 4,000–5,000 host UMIs per spot and 0–1,000 fungal UMIs per spot (**Figure 4b**). In contrast to the heart, no strong inverse relationship between host and fungal UMI distributions was observed in kidney tissue.

**Figure 4.**
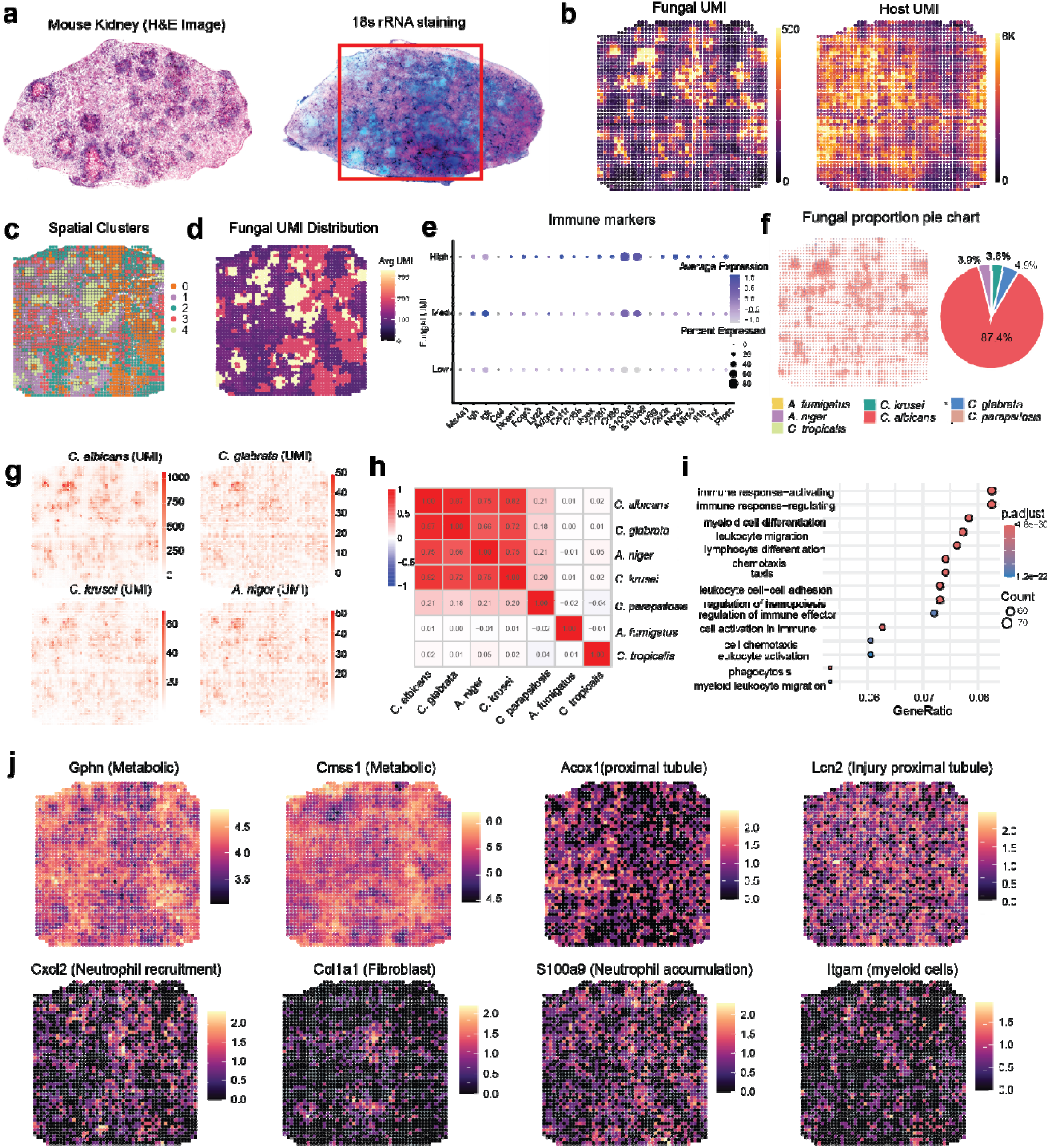
Spatially resolved host transcriptional programs in infected kindey associated with fungal burden revealed by fungal-DBiT. a, H&E staining of an infected heart section (left) and corresponding 18s rRNA staining image with the fungal-DBiT profiled region indicated (right). b, Spatial maps of fungal UMI counts and host UMI counts across the profiled region. c, Unsupervised spatial clustering of fungal-DBiT spots identifies distinct tissue niches within the infected region. d, Spatial distribution of fungal burden, highlighting infection-enriched microdomains. e, Dot plot summarizing differential expression of host genes across spots with high, medium and low fungal burden. f, Spatial pie-chart visualization and overall composition of fungal species across the profiled tissue. g, Spatial expression maps of representative fungal species, illustrating species-specific localization patterns. h, Pairwise correlation matrix of fungal species UMI abundance across spatial spots, revealing co-occurrence relationships among fungal taxa. i, GO analysis of fungal high burden spots. j, Spatial expression maps of representative host immune and inflammatory genes (including chemokines, cytokines, and myeloid markers), demonstrating localized host responses that co-localize with fungal-enriched regions.

We did unsupervised clustering and identified the major cell types in kidney (**Figure 4c, Figure S7a, s7b**). Cluster 0 and 4 show the highest fungal burden (**Figure S7d**). To examine how host immune programs vary with local fungal abundance, we stratified spatial spots by fungal UMI levels (Low, Medium, High) and evaluated the expression of canonical immune marker genes (**Figure 3e**). Spots with high fungal burden exhibited markedly increased expression of myeloid and innate immune markers, including *Ms4a1, Ighg1, Lgals3, Cxcl2,* and *Ccl7*, indicating localized recruitment or activation of inflammatory immune cells within fungal-enriched regions. In contrast, spots with low fungal burden showed minimal expression of these immune genes, consistent with a relatively quiescent immune microenvironment. Notably, genes associated with antigen presentation and leukocyte activation (*H2-D1, Cd80, Cd86*) were preferentially enriched in high–fungal UMI regions, suggesting active immune engagement at sites of fungal colonization. Intermediate fungal burden spots displayed a graded immune response, with moderate expression of chemokines and activation markers, supporting a continuum rather than a binary host response to fungal presence. Bulk RNA-seq was also conducted on the same kidney tissue and find similar fungal abundance as fungal-DBiT (**Figure S7e**).

Species-resolved spatial maps demonstrated that *C. albicans* is also the dominant fungal species in kidney tissue as in heart, accounting for approximately 87.4% of total fungal UMIs, with localized hotspots showing the highest abundance (**Figure 4f**). Analysis across spatial clusters showed consistent dominance of *C. albicans* within fungal-enriched regions, despite detectable co-occurrence of multiple species (**Figure 4j**). We show representative genes for different species and the dominant genes for *C. albicans* (**Figure S8**). Pairwise correlation and co-occurrence analyses revealed strong positive spatial associations among *Candida* species (Pearson’s r ≈ **0.7–0.9**), indicating differential niche overlap within the kidney microenvironment (**Figure 4h**). Integration of fungal abundance with host gene expression further identified localized associations between fungal-enriched regions and host inflammatory and stress-related genes, demonstrating the capacity of Fungal-DBiT to capture spatially resolved host–fungal interactions in renal tissue (**Figure 4i**).

Checking individual host genes reveals spatially organized host responses that integrate metabolic reprogramming, tissue injury, and immune infiltration within infected tissue (**Figure 4j and S8**). Metabolic genes (*Gphn*, *Cmss1*) show broad but structured expression patterns, indicating regional metabolic heterogeneity rather than uniform activation. In contrast, injury-associated markers (*Acox1*, *Lcn2*) are enriched in discrete areas, consistent with localized tissue damage. Immune-related genes further highlight focal inflammatory responses. *Cxcl2* and *S100a9* mark regions of active neutrophil recruitment and accumulation, while Itgam indicates localized myeloid cell infiltration. These immune-enriched areas partially overlap with, but are not identical to, fibroblast-associated regions marked by *Col1a1*, suggesting spatially distinct inflammatory and remodeling programs.

### Spatial profiling of low abundance fungi in mouse intestine

We applied Fungal-DBiT to Swiss-roll preparations of healthy mouse small intestine to assess baseline spatial distributions of fungal and host transcripts in the absence of overt infection (**Figure 5a**). Histological analysis revealed preserved intestinal architecture along the crypt–villus axis, and spatial transcriptomic profiling showed robust host RNA detection throughout the tissue, with comparatively low and heterogeneous fungal RNA signals (**Figure 5b**).

**Figure 5.**
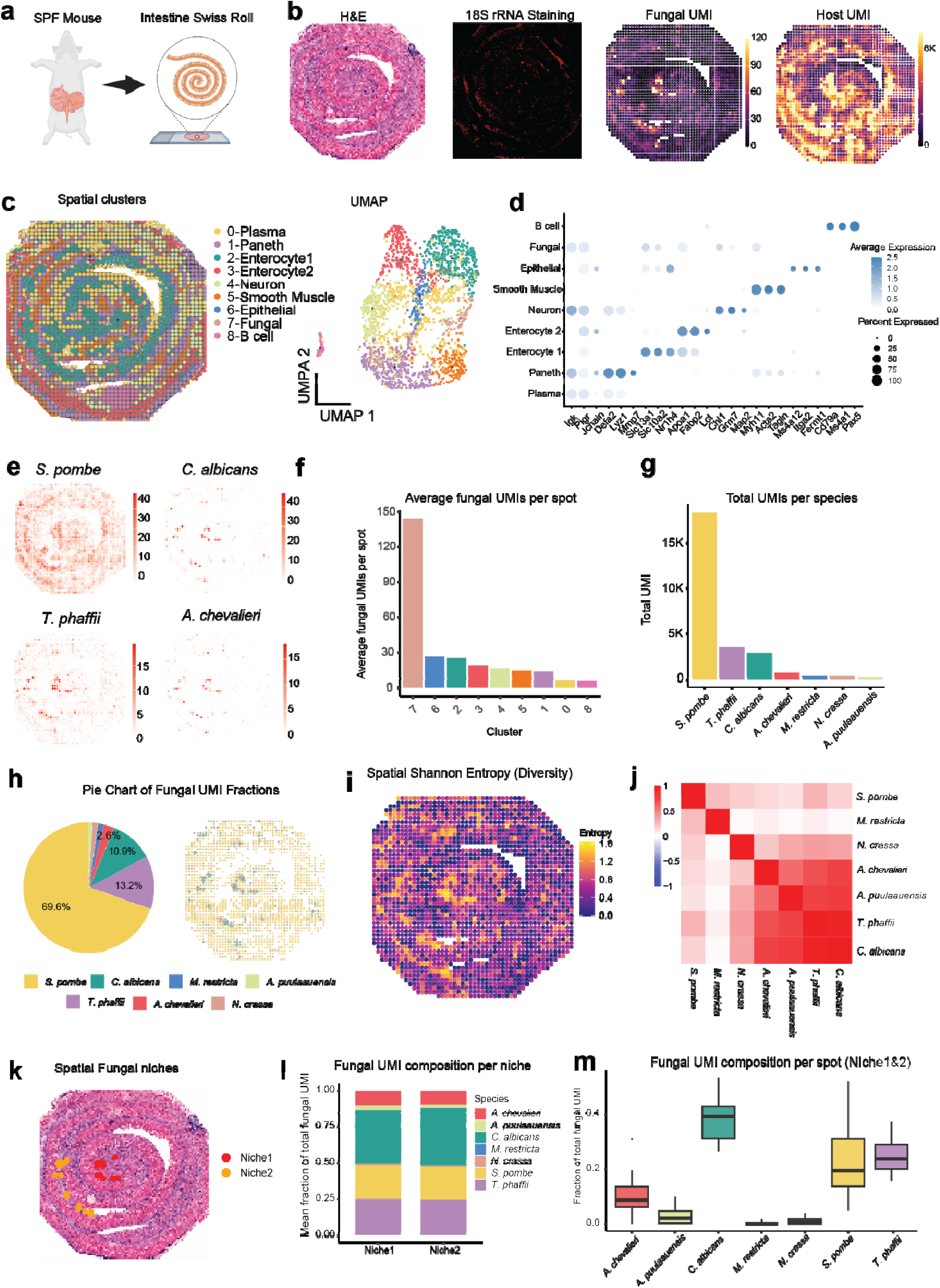
Spatial fungal niche organization in mouse intestine revealed by fungal-DBiT. a, Schematic illustrating the spiral sampling strategy and application of fungal-DBiT to an infected heart section. b, H&E staining of the tissue section and corresponding fungal 18S rRNA staining, confirming spatially patterned fungal localization within tissue architecture (left). Spatial maps of fungal UMI counts and host UMI counts, showing heterogeneous fungal burden across the spiral tissue section (right). c, Unsupervised spatial clustering of fungal-DBiT spots identifies multiple concentric tissue niches (left). UMAP embedding of spatial spots colored by cluster identity(right). f, Average fungal UMI counts per spatial cluster, highlighting clusters enriched for fungal transcripts. g, Total fungal UMI counts per species across the section. h, Pie chart and spatial map showing the fractional contribution of fungal species across the tissue. i, Spatial Shannon entropy map depicting local fungal diversity across the tissue section. j, Pairwise correlation matrix of fungal species abundance across spatial spots, revealing co-occurrence and exclusion patterns. k, Spatial identification of fungal niches overlaid on histology. l, Fungal species composition within distinct spatial niches. m, Distribution of fungal UMI counts per spot across niches, demonstrating niche-specific fungal load differences.

Unsupervised clustering identified multiple transcriptionally distinct domains corresponding to known intestinal structures, including epithelial, stromal, and muscle-associated regions (**Figure 5c**, **5d and S9**). Fungal UMIs were detected at low levels across clusters, with average fungal counts per spot markedly lower than those observed in infected tissues and restricted to scattered microdomains rather than contiguous infection cores (**Figure 5e**). Among these, cluster 7 represents a localized fungal-enriched niche.

Species-resolved spatial maps indicated that fungal signals in healthy intestine were dominated by a limited number of taxa, with *Saccharomyces*- and *Candida*-related signals detected at dominant abundance and shows specific niches forming (**Figure 5f and S10a**). *S. pombe* is a non-pathogenic yeast that can tolerate nutrient fluctuation and anaerobic conditions characteristic of the gut environment, making it well suited to persist in the intestine.^18^ In healthy mice, host immune surveillance tightly restricts the expansion of invasive fungi such as *C. albicans*, thereby limiting their contribution to the detectable fungal transcript pool. Global composition analysis showed that no single fungal species exceeded ∼70% of total fungal UMIs, and most species contributed small fractions, consistent with low fungal burden under physiological conditions (**Figure 5j**).

Spatial diversity analysis using Shannon entropy revealed relatively uniform and low fungal diversity across the tissue, without sharply demarcated niches of high fungal dominance (**Figure 5k**). Correlation and co-occurrence analyses showed modest positive associations among detected fungal taxa, but overall weaker correlations compared with infected tissues, reflecting sparse and dispersed fungal presence (**Figure 5i and 5m**).

Further analysis identified two discrete spatial fungal niches (Niche 1 and Niche 2) occupying distinct regions of the tissue section, indicating that fungal colonization is organized into localized microenvironments rather than being uniformly distributed (**Figure 5k and S11b**). Despite their spatial separation, the two niches exhibited highly similar fungal species composition, with *S. pombe* as a dominant component in both, accompanied by shared contributions from *C. albicans*, *A. pullulans*, and other low-abundance species (**Figure 5l**). Notably, per-spot analysis within each niche revealed considerable local variability in species dominance (**Figure 5m**), suggesting that while the overall community structure is conserved across niches, individual fungal microdomains can be driven by different taxa rather than a single uniformly dominant species.

## Discussion

Understanding host–microbiota interactions require technologies that can simultaneously profile host and microbial cells while preserving their native spatial organization. Although sequencing-based approaches have transformed our understanding of microbial communities, most existing methods lack spatial resolution, limiting insight into the tissue microenvironments where host–microbe interactions occur. This limitation is particularly pronounced for fungi, which are often low in abundance, structurally diverse, and spatially heterogeneous within tissues. In this study, we developed **Fungal-DBiT**, a microfluidic-based spatial transcriptomics platform that enables simultaneous, genome-wide profiling of host and fungal RNAs directly within intact tissue sections.

Using Fungal-DBiT, we spatially resolved fungal and host transcriptomes across multiple tissue contexts, including fungal-infected heart and kidney tissues as well as healthy small intestine. This approach allowed us to visualize localized fungal enrichment, delineate infection core and peripheral regions, and quantify species-resolved fungal distributions in situ. Importantly, Fungal-DBiT enabled integrative analysis of host transcriptional states within fungal-enriched niches, revealing spatially confined host responses associated with altered mitochondrial, contractile, and regulatory gene expression programs. These results demonstrate that fungal colonization is highly localized within tissues and is accompanied by distinct host transcriptional remodeling that would be obscured by bulk or non-spatial approaches.

From a technical perspective, Fungal-DBiT is scalable, cost-effective, and compatible with standard tissue processing workflows. The platform does not require physical separation or enrichment of fungal cells, preserving native host–fungal spatial relationships and minimizing experimental perturbation. Moreover, the ability to capture host and fungal transcriptomes within the same spatial framework enables direct interrogation of host–fungal interactions at genome-wide scale, providing a foundation for integrative transkingdom analyses. Despite these advances, several limitations remain. The current spatial resolution aggregates transcripts from multiple fungal and host cells within each capture spot, precluding single-fungus resolution. Future improvements in microfluidic chip design, spot density, and capture efficiency will be required to address these challenges. Integration with complementary modalities, such as spatial proteomics or targeted genomic assays, may further enhance resolution and mechanistic insight.

In summary, Fungal-DBiT represents, to our knowledge, the first microfluidic-based spatial transcriptomics platform capable of dual profiling genome-wide gene expressions from both host and fungal cells in situ. This work establishes a foundation for spatially resolved studies of fungal biology and host–fungal interactions and opens new avenues for investigating the roles of tissue-resident fungi in health, infection, and inflammatory disease.

## Methods

### Ethics

All animal procedures were approved by the Institutional Animal Care and Use Committee of Yale University. Mice were housed in standardized conditions at Yale University Animal Facility, and all experiments adhered to US legislative requirements.

### Mixed Invasive Fungal Infection Mouse Model

C57BL/6 mice, bred at the animal facility and gifts from Palm Lab, were treated with cortisone acetate (22.5 mg/mL, 10 μL/g) in 0.05% Tween-20 in PBS via subcutaneous injection every other day until the mice exhibited severe illness (e.g., >30% weight loss or lack of movement). A day after the first cortisone injection, mice were anesthetized using ketamine and administered a mixed fungal suspension containing 7 fungal strains (∼2 × 10^7^ CFU) derived from human clinical isolates, which had been stored at -80 °C prior to resuscitation. These strains were: *Candida parapsilosis*, *Candida glabrata*, *Candida albicans*, *Candida tropicalis*, *Aspergillus fumigatus*, *Aspergillus niger* and *Candida krusei*. The fungal suspension was applied under the tongue using cotton for oral inoculation. Hearts and kidneys were then harvested. Both organs were immediately embedded in OCT, flash-frozen on dry ice, and stored at −80 °C for subsequent experiments.

### SPF mouse model

Eight-week-old female C57BL/6J mice (Jackson Laboratory, stock no.000664) were harvested from gender-matched littermates and prepared using the same procedure. For complete observations of the niches and structures, we applied Swiss-rolling techniques onto the small intestine before embedding them in OCT.

### Swiss-Roll Technique for Intestinal Tissue

To preserve the full-length architecture of the intestinal tract and to enable comprehensive spatial profiling across regions, we applied the Swiss-roll technique to the small intestine and colon prior to embedding in OCT. After harvesting the intestines and transferring them to a clean petri dish with 1x PBS, we removed any mesenteric tissues, including fat, identified and transected the cecum. Then, we transected the small intestine from the cecum and put the intestines in PBS. The intestinal segments were gently flushed using a syringe with a gavage needle containing 10 mL Bouin’s fixative (50% ethanol/5% acetic acid in water) or cold 1x PBS to remove luminal contents. Tissues were cut open along the long longitude and briefly rinsed in 1x PBS. The proximal end of each segment was pulled toward the edge of a petri dish with the luminal side facing up. While holding the proximal end with forceps, the tissue was carefully rolled from the proximal to the distal end around a toothpick. Sterile forceps were used to pinch the wrapped edge against the toothpick to create a tight spiral. The small intestine were gently and gradually rolled to form a Swiss roll. The rolled tissue was transferred into a cryomold and embedded in OCT with the roll axis perpendicular to the bottom of the mold. This orientation allowed full cross-sectional visualization of the entire intestinal length in a single section. Embedded blocks were stored at −80 °C for downstream spatial transcriptomic and histological analyses.

### Tissue Sectioning

For fungal-DBiT experiments and corresponding histology staining, we sectioned the tissue into 10 μm thickness. Tissue sections were mounted on the center of poly-L-lysine-coated (PLL) slides (Electron Microscopy Sciences, 63478-AS) and stored at −80 °C for subsequent experiments.

### Microfluidic Device Fabrication

We fabricated microfluidic device using soft lithography as described previously^16^. Chrome photomasks with the specified channel widths were designed and ordered from Front Range Photomasks Inc. (Lake Havasu City, AZ). Master molds were fabricated in-house on clean silicon wafers using SU-8 negative photoresists (SU-8 2010, SU-8 2025, and SU-8 2050; MicroChem). Each photoresist was chosen to produce the desired channel height: ∼28 μm for 25-μm-wide channels (SU-8 2025), and ∼50 μm for 50-μm-wide channels (SU-8 2050). The photoresist was spin-coated following the manufacturer’s protocols and processed using standard photolithography, including soft baking, UV exposure through the chrome photomask, post-exposure baking, development, and hard baking, to generate the final SU-8 patterned molds.

To cast the PDMS chips, the elastomer base and curing agent were mixed at a 10:1 ratio by weight. Approximately 15 g of the mixture was poured into SU-8 molds when reusing molds, or 50 g was used for new molds cast in 10 cm Petri dishes to achieve a consistent PDMS thickness of ∼4 mm. The mixture was degassed under vacuum to remove air bubbles, then cured at 60 °C for 2 hours. Once cured, the PDMS slabs were carefully cut and peeled from the molds. Each chip was trimmed to fit standard tissue slide dimensions, and holes (∼2 mm diameter) were punched into the waffle manifolds to serve as fluid inlets and outlets for barcode delivery.

### Bulk RNA-seq

Total RNA was extracted from adjacent fungal infected sections using RNeasy Mini Kit (Qiagen). cDNA was generated using the SMART-Seq® v4 Ultra Low Input RNA Kit. Libraries were prepared from normalized RNA inputs ranging from 200 pg to 50 ng using the NEBNext Single Cell/Low Input RNA Library Prep Kit for Illumina (E6420L). Briefly, cDNA synthesis was performed using a template-switching approach, followed by fragmentation, end repair, dA-tailing, adapter ligation, and PCR amplification using the NEBNext Ultra II FS workflow. Indexed libraries that met predefined quality and quantity thresholds were quantified by quantitative PCR using a commercially available KAPA Biosystems kit. Library insert size distributions were assessed using the LabChip GX system or Agilent Bioanalyzer. Samples with a final library concentration of ≥0.5 ng/µL were advanced for sequencing. Sequencing was performed on an Illumina NovaSeq 6000 system at the Yale Center for Genome Analysis. RNA-seq reads were aligned to the appropriate reference genome using STAR.

### Spatial Host-Fungal RNA Sequencing

To preserve RNA integrity and spatial resolution during sample preparation, fresh-frozen tissue sections were equilibrated to room temperature for 10 min in a closed container. A PDMS reservoir was cut to fit the tissue region and set aside for later use. The tissue was then gently dried using an air pump. Fixation was performed by applying 200 μL of 4% paraformaldehyde (Electron Microscopy Sciences Inc) over the tissue, followed by a 20-minute incubation at room temperature. After fixation, the slide was washed twice with PBS containing 0.1 U/μL RNase inhibitor (RI). To quench residual aldehydes, 200 μL of 1.25 M glycine containing RI was added and incubated for 5 min, followed by two additional PBS-RI washes.

To enzymatically remove the fungal cell wall, 50 μL of freshly prepared digestion solution was applied directly to each tissue section. The digestion reagent consisted of 5 μL Zymolyase (5 U/μL, Zymo Research Corporation, cat# E1005), 5 μL dithiothreitol (100 mM, Thermofisher Scientific, cat# R0861), 0.5 μL Triton X-100 (10%), 1.2 μL RNase inhibitor (RI), and 38.3 μL nuclease-free water. Following application, the slide was incubated at 37 °C for 15 min to facilitate cell wall digestion and permeabilization. After incubation, the tissue section was gently washed twice with 0.5× PBS supplemented with RNase inhibitor (PBS-RI) to remove residual enzymes and digestion reagents.

Tissue permeabilization was carried out by applying 50 μL of 0.5% Triton X-100 in PBS directly onto the tissue section and incubating for 20 min at room temperature. This step facilitates membrane disruption and enhances reagent accessibility for subsequent reactions. Following permeabilization, the slides were washed twice with PBS supplemented with RNase inhibitor to remove residual detergent. The slides were then briefly dipped in nuclease-free water to eliminate salt and were dried using a gentle stream of air before proceeding to downstream processing steps.

To initiate reverse transcription, a 60 μL reverse transcription master mix was freshly prepared immediately before use. The RT mix consisted of 12 μL 5× RT buffer, 15 μL 30% PEG8000, 11 μL nuclease-free water, 0.4 μL RNase inhibitor, 0.6 μL Superase-In RNase inhibitor, 3 μL dNTPs (10 mM each), 6 μL Maxima H Minus Reverse Transcriptase (ThermoFisher), and 12 μL RT primer (25 μM). The prepared RT master mix was carefully dispensed onto the tissue region of interest (5 × 5 mm) within a PDMS reservoir affixed to the slide. The reservoir was sealed with parafilm to prevent evaporation, and reverse transcription was performed by incubating the slide for 30 min at room temperature to allow primer annealing, followed by 90 min at 42 °C in a humidified chamber to enable cDNA synthesis.

After reverse transcription, slides were washed with 1× PBS for 5 min, dipped in water three times, and dried using an air pump. First ligation was performed by loading 5 μL of barcode A ligation mix to each of the 50 inlets of a PDMS chip. The barcode A ligation mix was freshly prepared by combining 68 μL nuclease-free water, 29 μL 10× T4 DNA ligase buffer (New England Biolabs), 11 μL T4 DNA ligase (400 U/μL, New England Biolabs), 2 μL RNase inhibitor (40 U/μL, Enzymatics), 0.7 μL Superase-In RNase inhibitor (20 U/μL, Ambion), 5.4 μL 5% Triton X-100, and 116 μL 1× NEBuffer 3.1 containing 1% RNase inhibitor. The barcode mixture was vacuum driven through the microfluidic channels for 1 min, followed by incubation at 37 °C for 15 min. This process was repeated once to increase ligation efficiency. The PDMS chip was then removed, and a second chip was applied perpendicularly. Barcode B ligation was performed using the same process. The tissue images with PDMS chips were imaged before and after ligation.

For tissue lysis, 70 μL of lysis solution was prepared by combining 10 μL proteinase K (20 mg/mL; New England Biolabs) with a mixture of 1× PBS and 2× lysis buffer. The 2× lysis buffer consisted of 40 mM Tris-HCl (pH 8.0), 800 mM NaCl, 200 mM EDTA (pH 8.0), and 8.8% SDS. The lysis solution was applied to the tissue section and incubated in a humidified chamber at 55 °C for 2 h. Following incubation, the lysate was collected into a 1.5-mL DNA low-bind tube, adjusted to a final volume of 80–100 μL with lysis buffer, and further incubated overnight at 55 °C to ensure complete protein digestion.

On the following day, lysates were purified using the DNA Clean & Concentrator kit (Zymo Research) at a 1:7 sample-to-binding buffer ratio. DNA was eluted in 150 μL of nuclease-free water preheated to 70 °C, reapplied to the column, and collected into a DNA low-bind tube. Biotinylated cDNA was subsequently captured using 40 μL Dynabeads MyOne Streptavidin C1 beads (ThermoFisher) that had been pre-washed in 1× B&W buffer containing 0.05% Tween-20. The beads were incubated with the purified cDNA for 1 h at room temperature with rotation, followed by three washes with B&W buffer and 10 mM Tris-HCl (pH 8.0) supplemented with 0.1% Tween-20.

Template switching was carried out on bead-bound cDNA using a total reaction volume of 220 μL per sample. The reaction mix contained 44 μL 5× Maxima RT buffer, 51 μL nuclease-free water, 44 μL 30% PEG8000, 44 μL 20% Ficoll PM-400, 22 μL dNTPs (10 mM each), 5.5 μL Maxima H Minus Reverse Transcriptase (ThermoFisher), and 5.5 μL template-switching primer (100 μM). The beads were incubated with the reaction mixture for 30 min at room temperature, followed by 90 min at 42 °C to facilitate template switching. After incubation, the beads were washed twice with 10 mM Tris-HCl (pH 8.0) supplemented with 0.1% Tween-20 and twice with nuclease-free water.

PCR amplification was conducted in a 220 μL reaction using 110 μL 2× Kapa HiFi HotStart Master Mix (Roche), 8.8 μL of each 10 μM primer (Primer 1 and Primer 2), and 92.4 μL nuclease-free water. The reaction cycle was: 95 °C for 3 min; 5 cycles of 98 °C for 20 seconds, 65 °C for 45 seconds, and 72 °C for 3 min; followed by 72 °C for 5 min. Amplified cDNA was purified using a 0.7× SPRI bead ratio and eluted in 20 μL nuclease-free water.

Library preparation was carried out using the Nextera XT DNA Library Prep Kit (Illumina, FC-131-1024). A total of 1–2 ng purified cDNA was diluted to 5 μL in nuclease-free water and combined with 10 μL Tagment DNA buffer and 5 μL Amplicon Tagment Mix. The mix was incubated at 55°C for 5 min. After adding 5 μL NT buffer and incubating at room temperature for 5 min, 15 μL PCR master mix, 8 μL nuclease-free water, and 1 μL each of P5 and indexed P7 primers (10 μM) were added. The PCR program was 95°C for 30 seconds; 12 cycles of 95°C for 10 seconds, 55°C for 30 seconds, and 72°C for 30 seconds; followed by 72°C for 5 min. Final libraries were purified with 0.7× SPRIselect beads and eluted in 20 μL nuclease-free water. Library size (400–600 bp) was verified using an Agilent Tapestation.

Sequencing was performed on an Illumina NovaSeq Plus platform. For each tissue section, approximately 200 million sequencing reads were requested to ensure sufficient coverage for downstream analyses.

### Barcodes, Primers, and Oligos Design

All oligonucleotides were synthesized by IDT and purified by HPLC. Barcodes were designed to capture mRNA transcripts and encode two-dimensional spatial information through in situ ligation between orthogonal oligos (Barcode A and Barcode B). A separate reverse transcription (RT) primer, not part of the barcode sets, was synthesized with a 5′ phosphate and an internal biotin-dT modification for downstream cDNA capture via streptavidin purification.

Barcode A encodes row information and consists of either 50 unique oligos, each with a 5′ phosphate modification and three functional segments: (1) a poly(T)-VN region (15 nt + VN) that serves as a reverse transcription primer, (2) a ligation linker 1 region (∼22 nt) with a 14 bp overlap with linker 2 in Barcode B, and (3) an 8 nt position-specific spatial barcode embedded between the ligation linker and poly(T) region to encode row-specific identity.

Barcode B complements Barcode A, encoding column information. First, a ligation linker by 14 bp was introduced, followed by a 10 nt unique molecular identifier (UMI), a spatial barcode region (∼23 nt), and a PCR handle (13 nt). Barcode B oligos also contain a 5′ biotin modification for streptavidin-based cDNA capture. All Barcode B oligos are synthesized as single strands. Sets of 50 Barcode B sequences were paired with Barcode A to enable two-dimensional spatial mapping. All oligo sequences are listed in Tables S2-S4.

### Fluorescence in Situ Hybridization

A general fungal fluorescent labelled 18S rRNA targeting probe (5′-Cy5-GTGATGCCCTTAGATGT-3) was used, enabling detection of a broad range of fungal taxa. To perform the experiment, tissue sections were fixed by 200 μL of 4% paraformaldehyde for 15 min at room temperature, followed by three washes in 1× PBS and complete air drying. The probe was first thawed to room temperature while the hybridization buffer was prewarmed to 50°C to facilitate probe-target annealing. To facilitate probe penetration, fungal cell walls were enzymatically permeabilized prior to 18S rRNA FISH by Zymolyase. Permeabilization was performed by incubating slides in 200 μL of 0.5% Triton X-100 for 20 min at room temperature to increase probe access. Slides were rinsed briefly with PBS before applying 20 μL of FISH probe at a final concentration of 10 ng/μL. Hybridization was carried out in a humidified chamber for 2–3 h or overnight in the dark to minimize nonspecific binding and photobleaching.

After hybridization, slides were washed once in prewarmed wash buffer for 10 min, followed by two additional washes at room temperature. Slides were soaked briefly in distilled water to remove residual salts. Nuclear counterstaining was performed using Hoechst dye diluted at 2 drops/mL in PBS, incubated on the slides for 20 min at room temperature, and protected from light. After three washes in PBS, the slides were fully air dried. To preserve signal and reduce photobleaching, 50 μL of Vectashield antifade mounting medium (cat# ZK0303, Vector Laboratories) was applied. Coverslips were gently placed over the tissue, and slides were stored at 4 °C until imaging.

### Hematoxylin and eosin (H&E) Staining

The slides were fixed using the same protocol as the spatial sections and submitted to the Yale Department of Pathology, where H&E staining was performed using standard histological procedures. After staining, whole-tissue imaging was carried out using tile scanning to capture the complete architecture for morphological reference.

### Data processing

For sequence alignment and generation of the gene expression matrix, RNA data mapping followed our previous report.^16^ Briefly, Read 1 was processed using the ST pipeline v.1.8.1, mapping against combined mouse genome (GRCm38) and genome for individual bacteria species, while Read 2 was processed for extraction of unique molecular identifiers (UMIs) and spatial Barcode A and Barcode B. Spatial barcodes corresponding to microfluidic channels were demultiplexed, and the ENSEMBL ID matrix was converted to a gene names matrix for downstream analysis. The identification of pixel locations on tissues was performed using customized matlab scripts using MATLAB 2023B.

### RNA data clustering and visualization

For data normalization and variance stabilization, we applied the SCTransformV2 method (v0.3.5) implemented in Seurat (v5.0). Because fungal transcripts are spatially sparse and highly heterogeneous across tissue locations, a modified normalization strategy was used in which host genes served as reference features. Specifically, fungal genes were excluded from the reference feature set using the *residual.features* parameter in SCTransformV2, while remaining included in the final normalized expression matrix. This approach prevents fungal abundance from influencing the estimation of normalization parameters while preserving fungal signal in the normalized data.

For quantitative visualization, total raw UMI counts per fungal species were used to generate violin plots, summary pie charts, and bar plots. Total normalized UMI counts per fungal species were used for spatial pie-chart visualizations. In spatial pie charts, the proportion shown for each pixel corresponds to the square-root– transformed total normalized UMIs for each fungal species to improve visualization across a wide dynamic range of abundance. Pixel size was determined by thresholding the summed normalized fungal signal. Only fungal species with substantial cumulative abundance across all spatial locations were included, and spatial charts were displayed only at pixels exceeding a minimum fungal abundance threshold. Spatial co-localization was assessed by calculating Pearson correlation coefficients between total normalized RNA counts of fungal species across fungus-enriched pixels and visualized as co-localization heatmaps.

For tissue region annotation, paired samples were concatenated after excluding fungal genes. Dimensionality reduction was performed on the concatenated normalized expression matrix using principal component analysis (PCA) with the top 10 components. Batch correction was applied to the PCA embeddings using Harmony (Harmonypy v0.0.5), followed by neighborhood graph construction using the default implementation in Scanpy (v1.9.1). Tissue regions were identified using Leiden unsupervised clustering, and UMAP visualizations were generated based on the constructed neighborhood graphs with default Scanpy parameters. Differentially expressed genes (DEGs) defining each cluster were identified using the *FindMarkers* function for pairwise comparisons between pixel groups, and expression heatmaps were generated to display the top five DEGs per cluster.

### KEGG pathway enrichment analysis

The common targets were used for GO and KEGG pathway enrichment with the Cluster Profiler package in R software (v3.6.3), and the “ggplot2” package was used to visualize the GO and KEGG enrichment results.

### Metagenomics analysis using Kraken2

For host-associated mycobiome analysis, RNA-sequencing reads were first aligned to the mouse reference genome using STAR to identify host-derived transcripts. Reads mapping to the mouse genome were removed from the original FASTQ files to minimize host contamination. The remaining unmapped reads, enriched for fungal sequences, were extracted and compiled into new FASTQ files for downstream mycobiome profiling and taxonomic classification using Kraken2.^19^

For taxonomic classification, the Kraken2 Standard Database was initially used to identify fungal taxa and estimate their relative abundances. Based on these results, fungal species with the highest read counts were selected for further analysis. Reference genome files for these high-abundance fungal species were manually downloaded from NCBI and combined into a custom fungal reference. The fungal-specific FASTQ files were then processed using the ST pipeline (v1.8.1) to generate gene-level expression matrices for downstream analyses.

## Data availability

The sequencing data will be available on request. The original sequencing data and processed expression matrix data have been deposited to NCBI Gene Expression Omnibus (GEO) with access number GSE316403. The reviewer access token is: czgdwukqtrcdjaj

## Code availability

The python and R codes for data analysis and visualization are shared through github: https://github.com/edicliuyang/Fungal-DBiT

## Acknowledgements

The research is supported by Maximizing Investigators Research Award (MIRA) for Early-Stage Investigators 1R35GM150838 (to Y. L.). The research is also supported, in part, by Grant #IRG-21-132-60-IRG from the American Cancer Society. We also acknowledge the support of Leslie H. Warner Postdoctoral Fellowship to D. W.

## Author Contributions

X.Y. and Y.L. designed the study. X.Y. and F.G. performed the fungal-DBiT experiments. X.Y. and Y.L. conducted the data analysis. D.S. designed the fungal community, generated the mouse models and prepared mouse specimens. G.L., S.W., D.W., X.D., L.L., J.X., D.Z., Y.J., and J.D. contributed to chip fabrication and tissue sectioning. X.Y. and Y.L. wrote the initial manuscript. N.W.P. and Y.L. supervised the project.

