## Supplemental information for "Spatially resolved host–fungal transcriptomics profiling by fungal-DBiT"

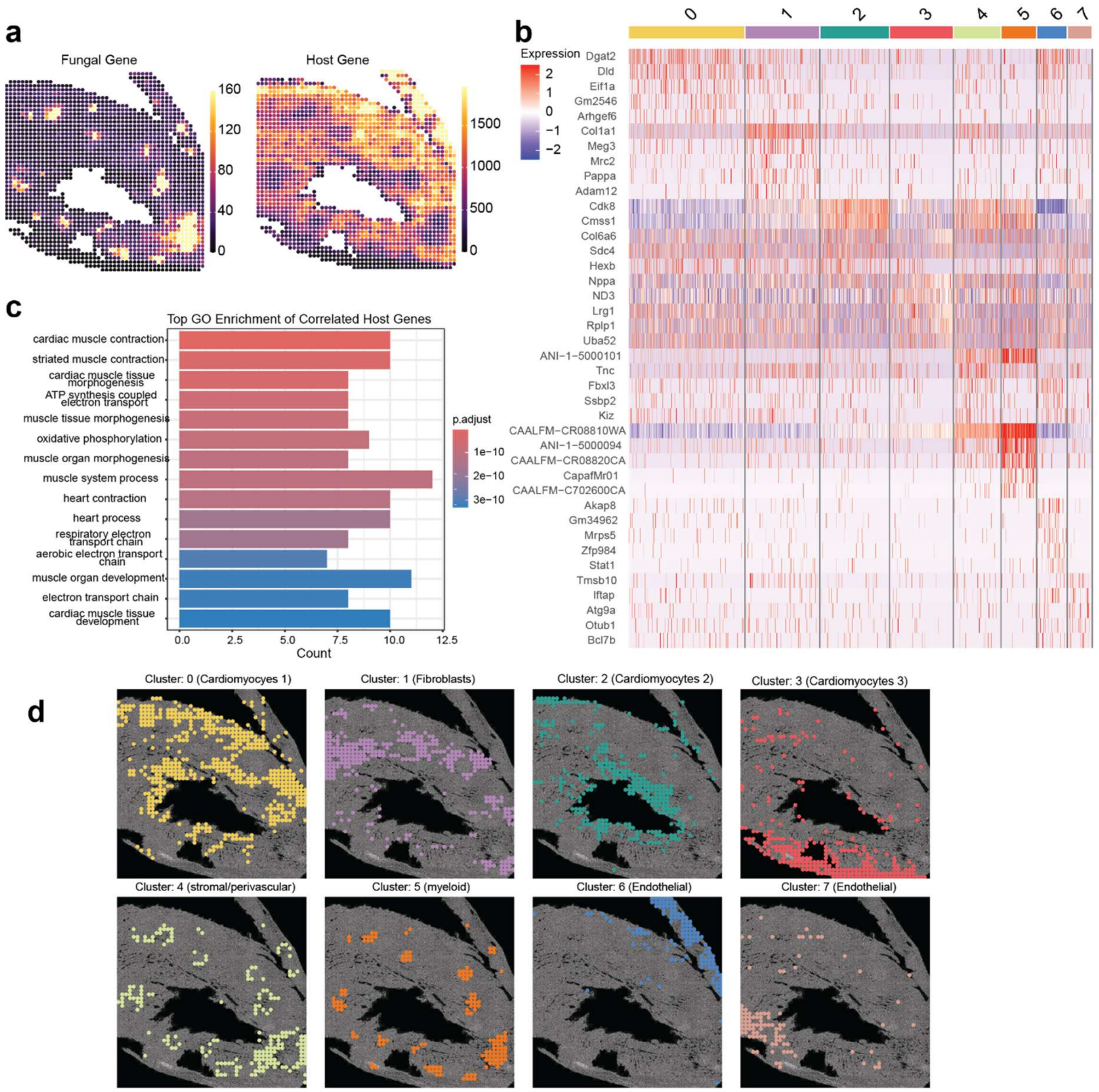


**Figure S1. Spatial host cell organization and transcriptional programs in heart tissue**. (a) Spatial maps of fungal and host UMIs. (b) Heatmap of marker gene expression across spatial clusters with cluster identities annotated. (c) Gene Ontology enrichment of cluster 0 host marker genes. (d) Spatial distribution of eight transcriptionally defined clusters, revealing cardiomyocyte, fibroblast, myeloid, stromal/perivascular, and endothelial cells.

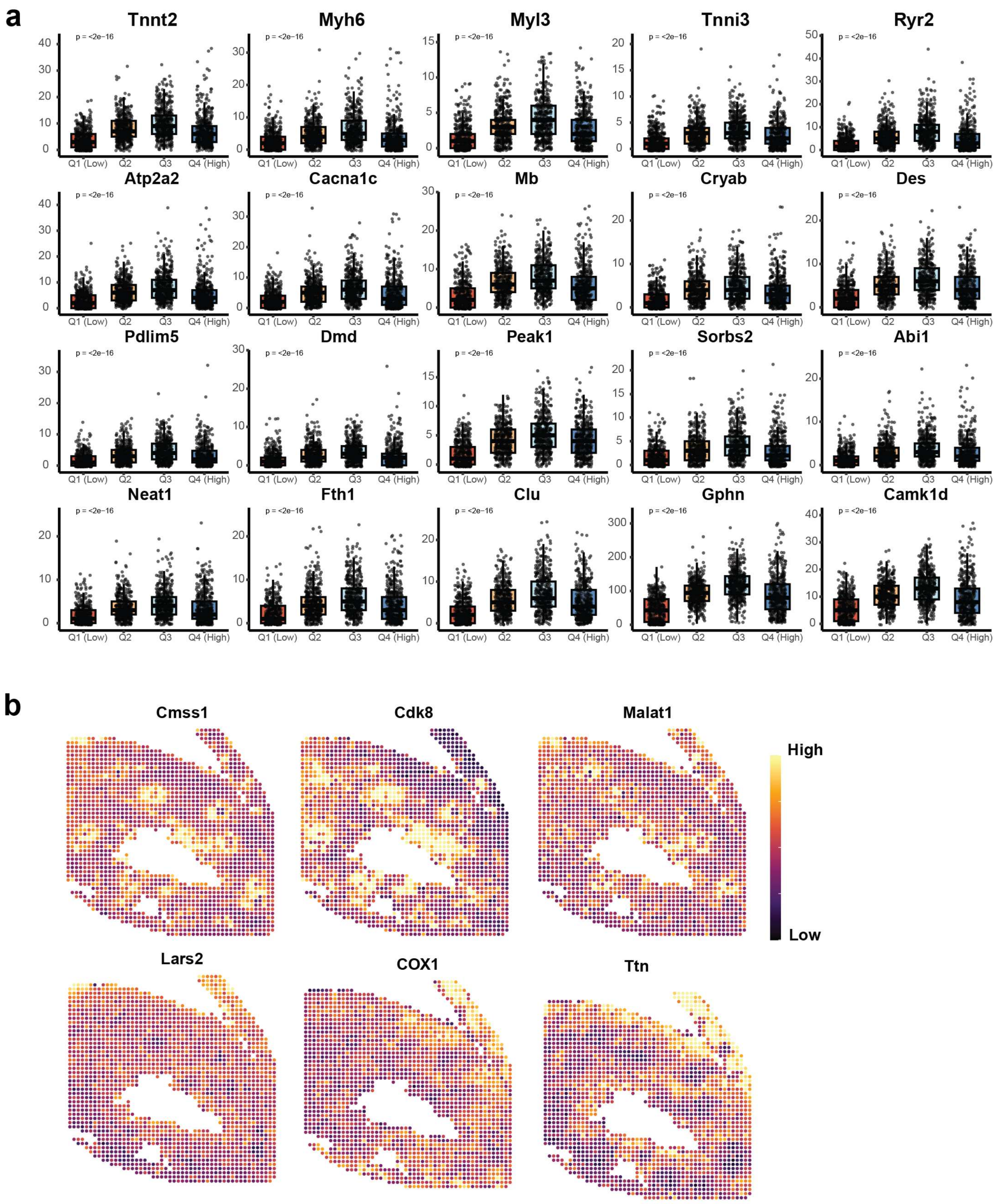


**Figure S2. Host gene analysis.** (a) Differential host gene expression between fungal infection core and periphery regions. (b) Spatial expression maps of representative genes highlight cluster-specific transcriptional patterns.

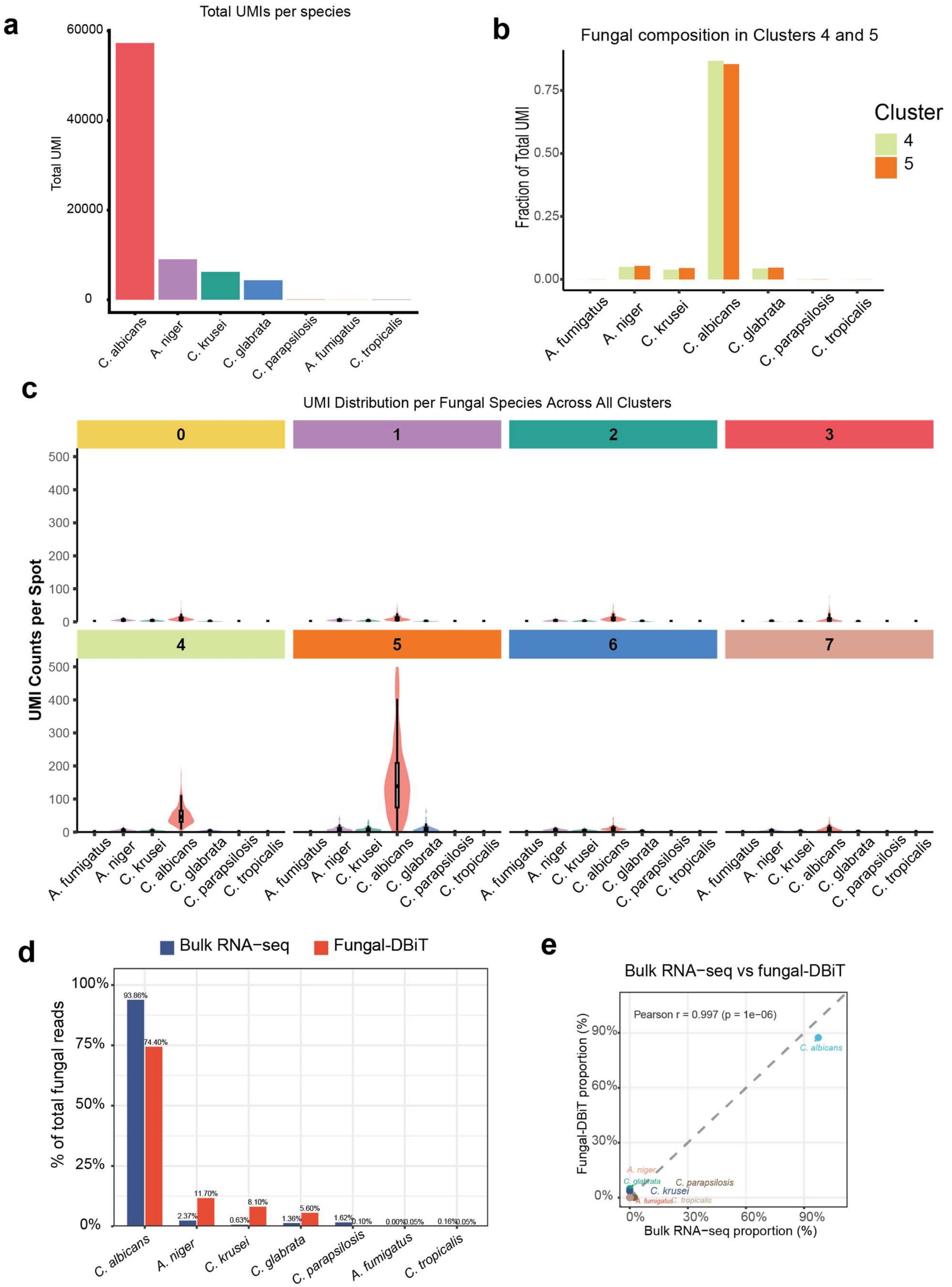


**Figure S3. Spatial analysis of host-fungal transcriptomics in mouse heart.** (a) Total fungal UMI counts per species (b) fungal composition in enriched clusters 4 and 5 (c) fungal UMI in each of the 8 clusters. (d) comparison between bulk RNA-seq and fungal-DBiT sequencing results. (e) correlation analysis.

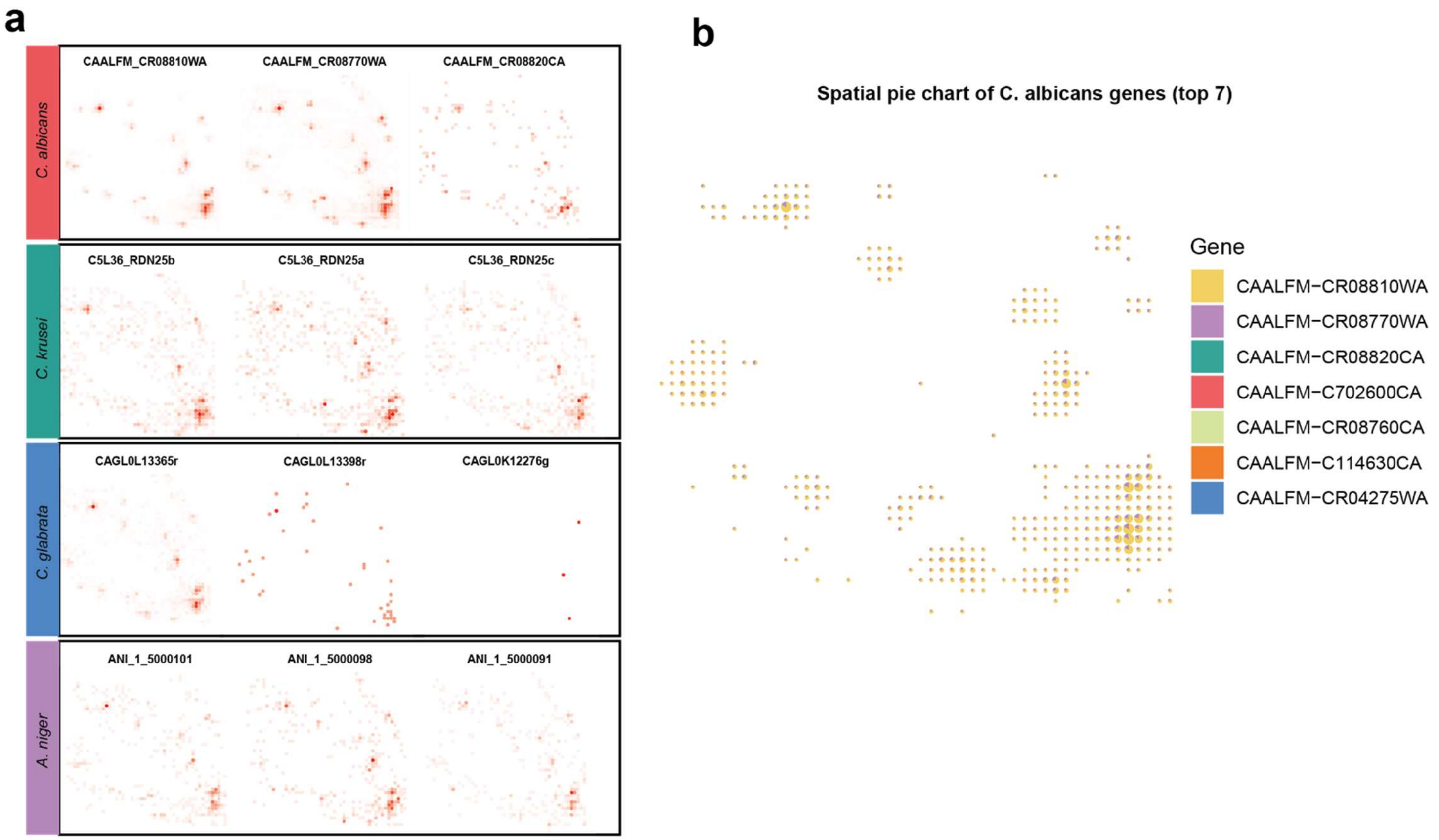


**Figure S4. Spatial distribution and gene-level composition of fungal transcripts.** (a) Spatial maps of representative fungal genes showing distinct localization patterns across the tissue section. (b) Spatial pie-chart visualization of the top expressed C. albicans genes, illustrating heterogeneous gene-level composition within fungal-enriched regions.

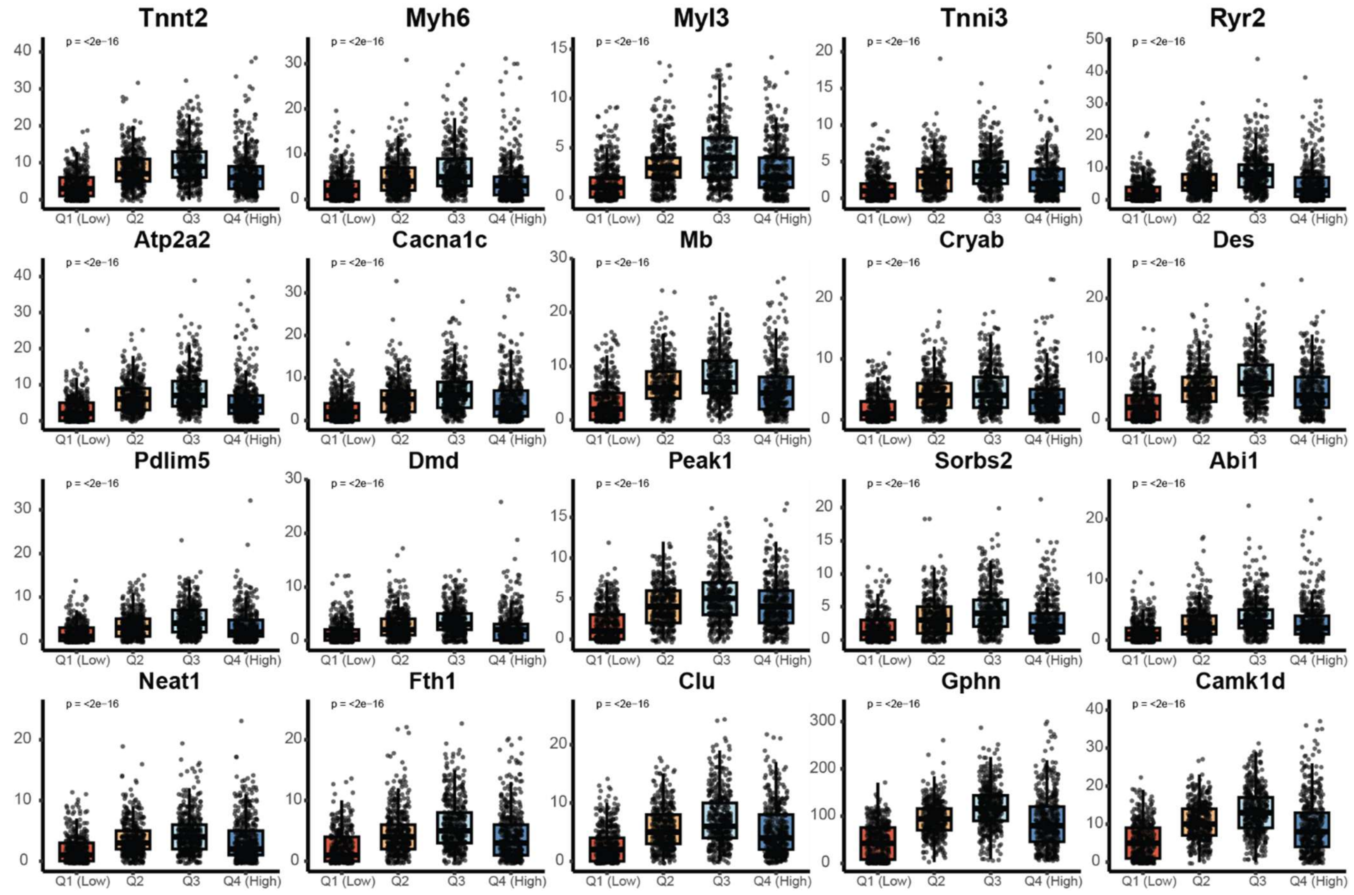


**Figure S5. Box-and-jitter plots of cardiomyocyte and stress-response genes across quartiles of fungal burden (Q1–Q4).**

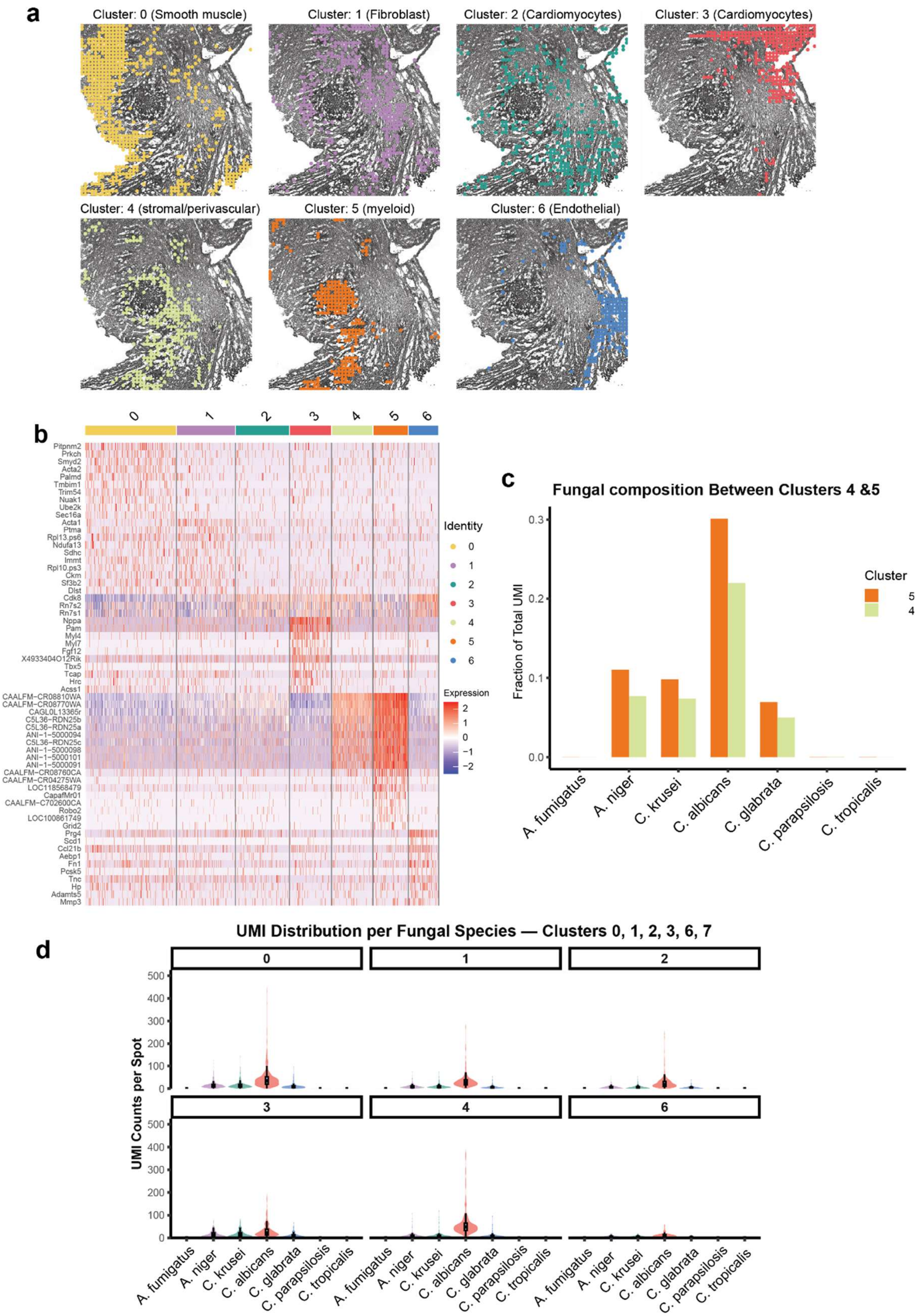


**Figure S6. Cluster-resolved fungal burden and spatial cell-type organization.** (a) Spatial maps of transcriptionally defined clusters, revealing distinct smooth muscle, fibroblast, cardiomyocyte, stromal/perivascular, myeloid, and endothelial regions. (b) Heatmap of cluster-specific marker gene expression. (c) Relative fungal species composition in clusters 4 and 5. (d) Violin plots showing fungal UMI distributions by fungal species across spatial clusters.

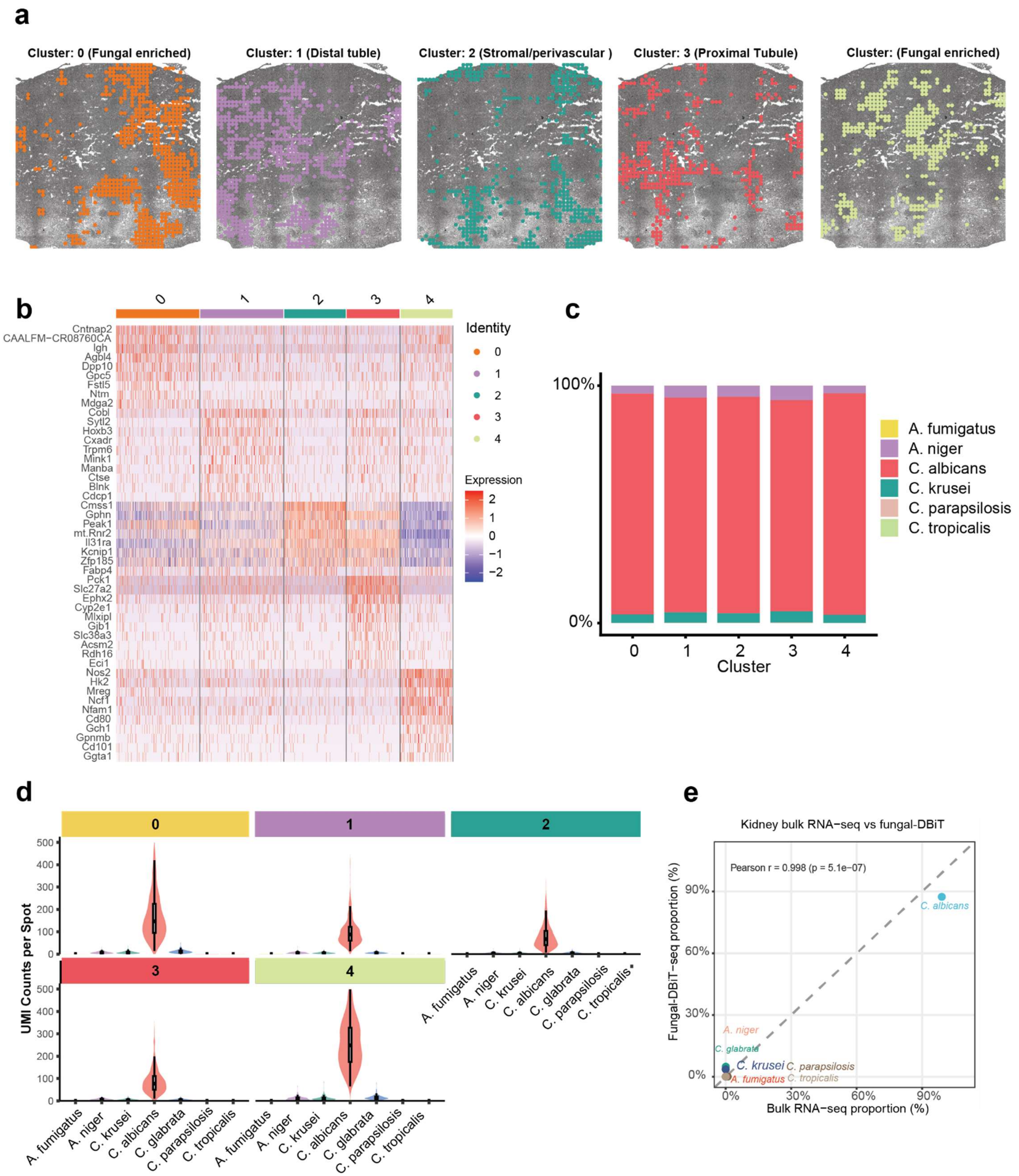


**Figure S7. Spatial clustering and associated fungal distributions.** (a) Spatial maps of five transcriptionally defined clusters overlaid on tissue morphology. (b) Heatmap of cluster-specific marker gene expression. (c) Relative fungal species composition across clusters. (d) Violin plots showing fungal UMI distributions across spatial clusters. (e) correlation analysis of bulk RNA-seq and fungal-DBiT results.

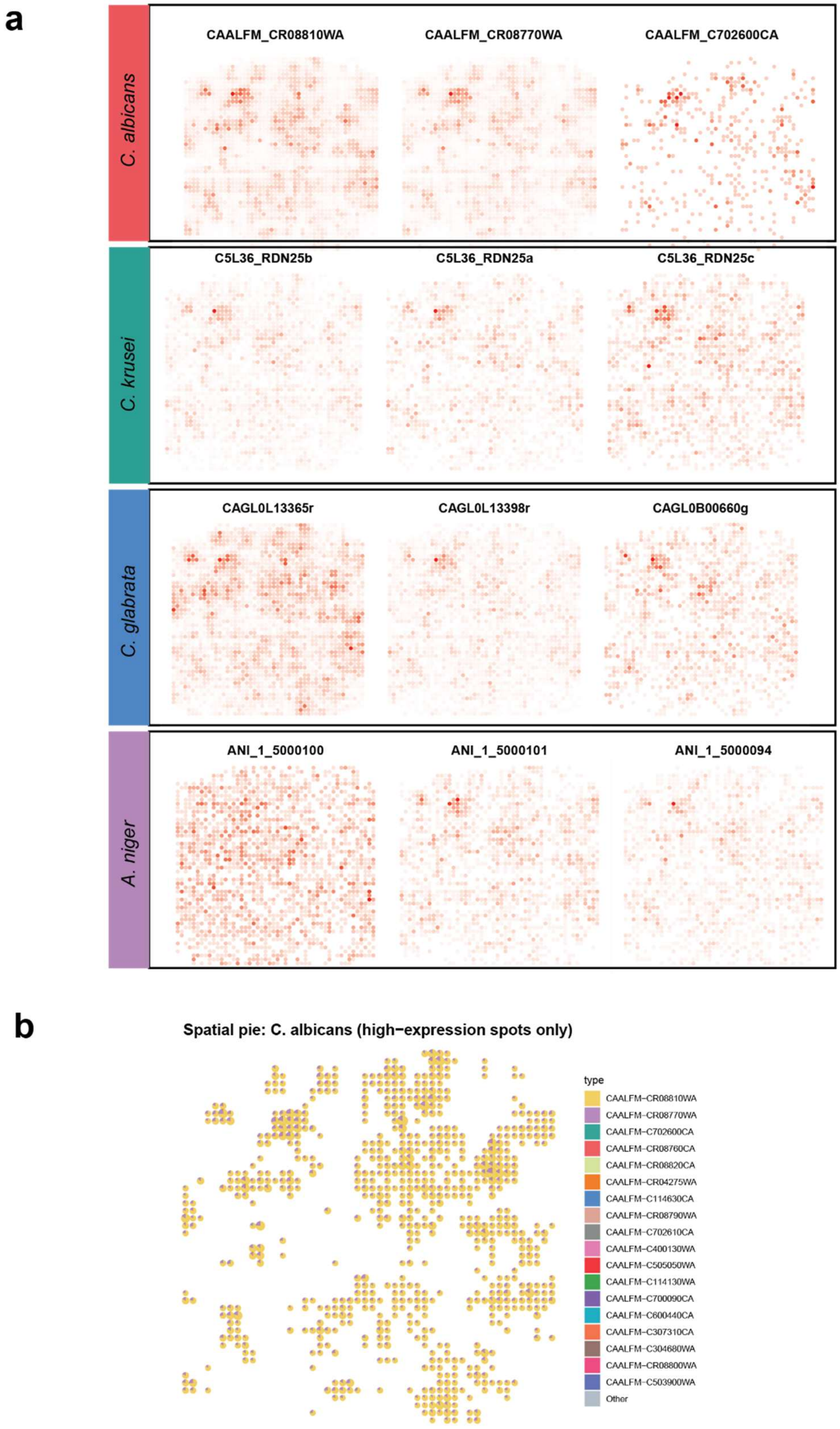


**Figure S8. Gene level spatial plot in mouse kidney.** (a) Spatial expression maps of representative genes from C. albicans, C. krusei, C. glabrata, and A. niger across the tissue section. (b) Spatial pie-chart visualization of C. albicans gene composition in high-expression spots, revealing heterogeneous gene usage within fungal-enriched regions.

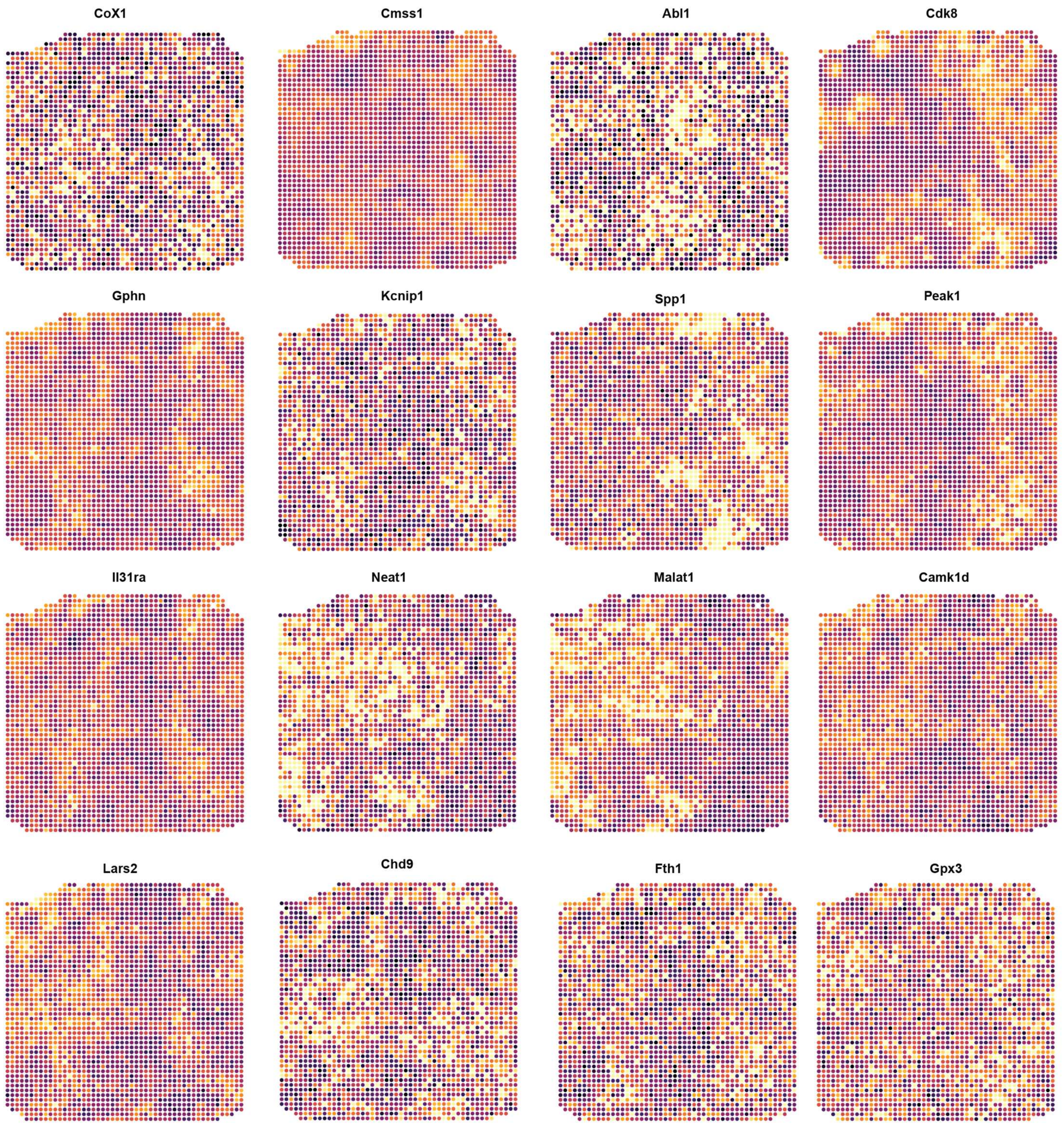


**Figure S9. Spatial expression patterns of representative host genes.** Spatial heatmaps of selected host genes across the tissue section, illustrating heterogeneous transcriptional activity and preserved spatial organization.

**a**

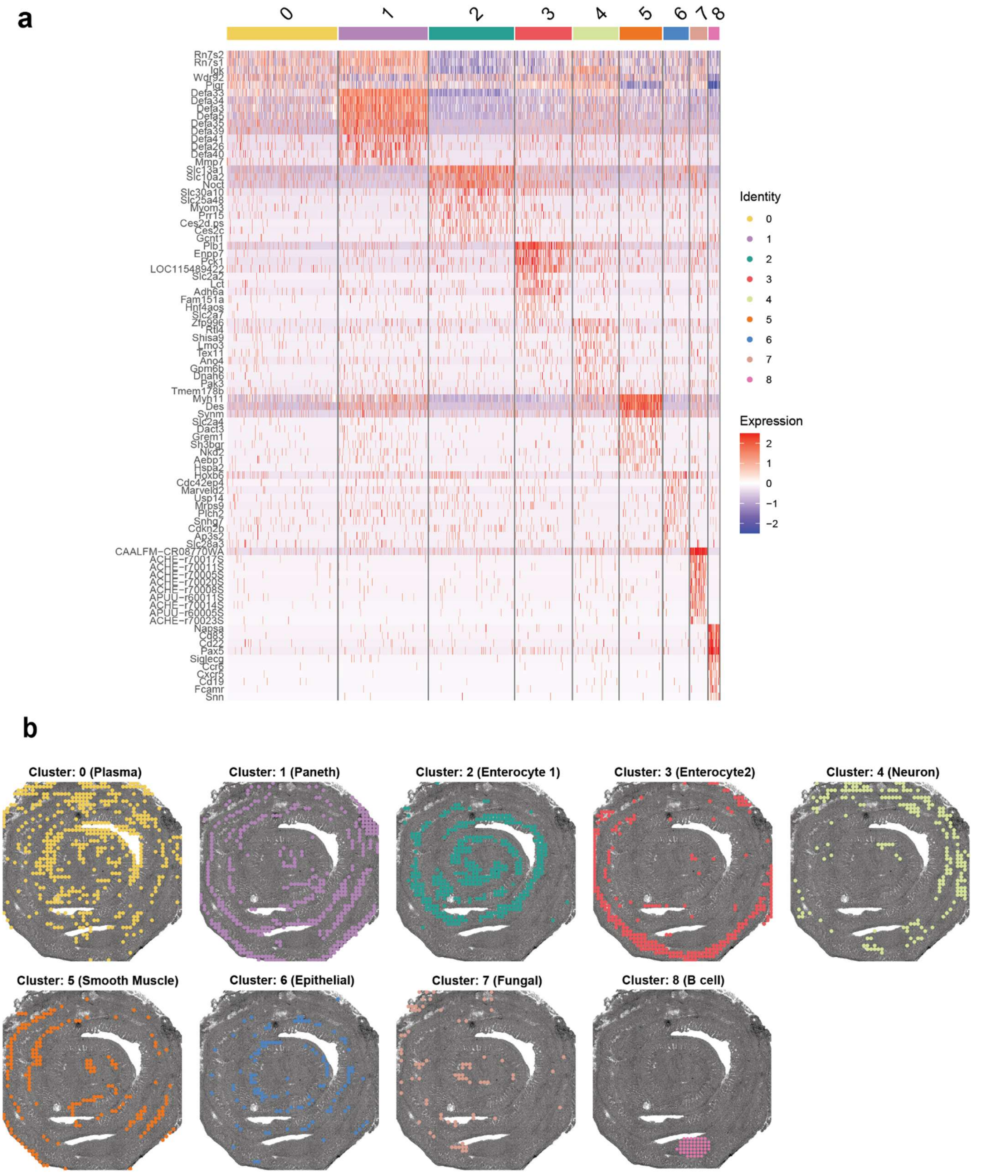


**b**



**Figure S10. Spatial clustering identifies host cell types and fungal-enriched regions in mouse small intestine.** (a) Heatmap of marker gene expression across spatial clusters with cluster identities indicated. (b) Spatial maps of clusters overlaid on tissue morphology, revealing plasma, Paneth, enterocyte, neuronal, smooth muscle, epithelial, and fungal-associated niches.

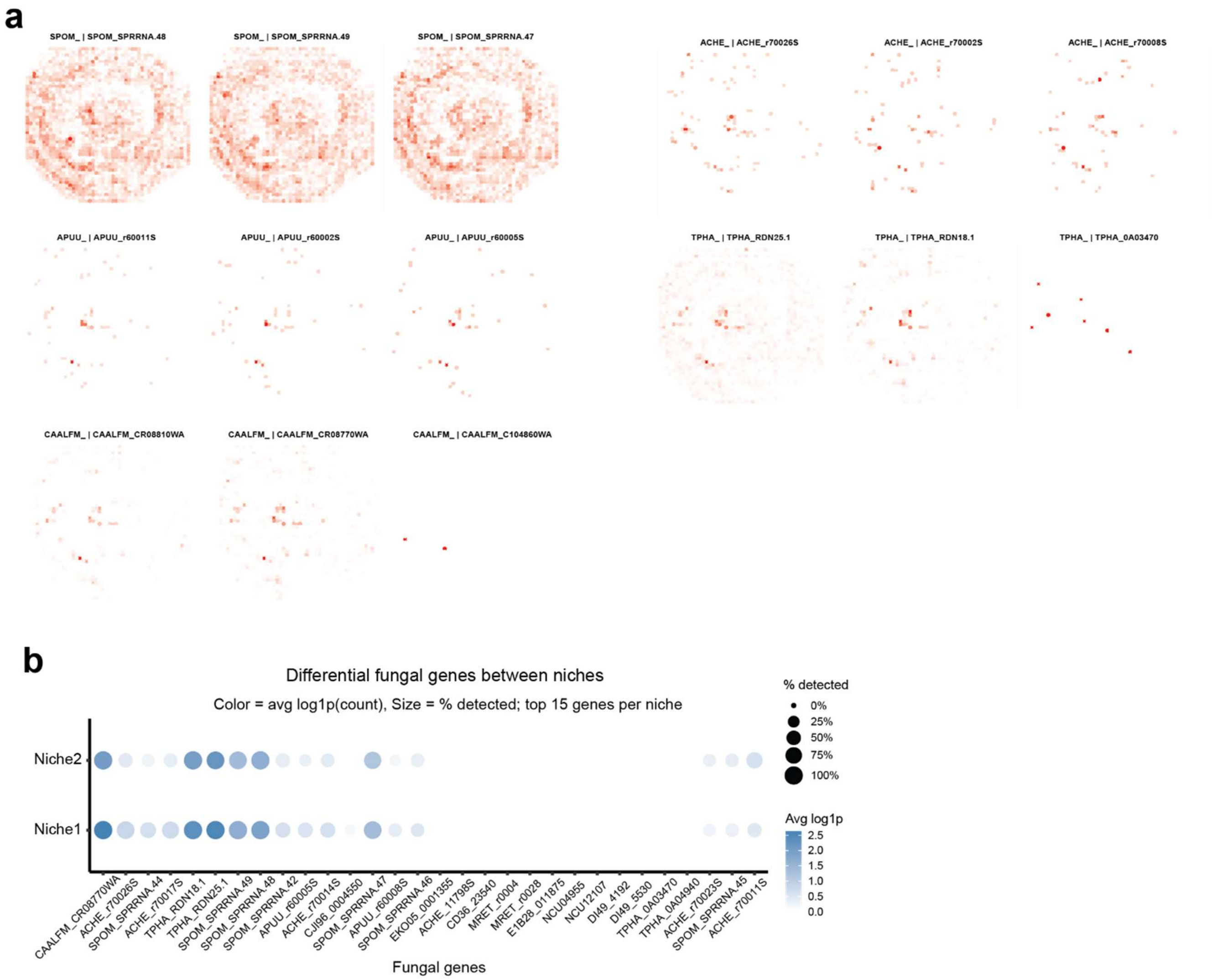


**Figure S11 | Spatial heterogeneity of fungal gene expression across tissue niches.** (a) Spatial maps of representative fungal genes showing heterogeneous and sparse localization patterns across the tissue section. (c) Dot plot of differentially expressed fungal genes between spatial niches, summarizing relative expression levels and detection frequencies.

**Table S1. Fungal species used in current research**

| Family | Species | Strain / isolate |
|---|---|---|
| Aspergillaceae | *Aspergillus fumigatus* | Af293 |
| Aspergillaceae | *Aspergillus niger* | CBS 513.88 |
| Debaryomycetaceae | *Candida albicans* | SC5314 |
| Saccharomycetaceae | *Candida glabrata* / *Nakaseomyces glabratus* | CBS138 |
| Pichiaceae | *Candida krusei* / *Pichia kudriavzevii* | Unspecified |
| Debaryomycetaceae | *Candida parapsilosis* | CDC317 |
| Debaryomycetaceae | *Candida tropicalis* | MYA-3404 |

**Table S2. DNA oligos used in fungal-DBiT**

| Sequence Name | Sequence (5'-3') |
|---|---|
| RT primer | /5Phos/CATCGGCGTACGACTNNNNNNNNNN/iBiodT/TTTTTTTTTTTTTTTVN |
| Template switch primer | AAGCAGTGGTATCAACGCAGAGTGAATrGrG+G |
| Ligation linker 1 | AGTCGTACGCCGATGCGAAACATCGGCCAC |
| Ligation linker 2 | CGAATGCTCTGGCCTCTCAAGCACGTGGAT |
| PCR Primer 1 | CAAGCGTTGGCTTCTCGCATCT |
| PCR Primer 2 | AAGCAGTGGTATCAACGCAGAGT |
| N501 | AATGATACGGCGACCACCGAGATCTACACTAGATCGCTCGTCGGCAGCGTCAG ATGTGTATAAGAGACAG |
| N502 | AATGATACGGCGACCACCGAGATCTACACCTCTCTATTCGTCGGCAGCGTCAGA TGTGTATAAGAGACAG |
| N701 | CAAGCAGAAGACGGCATACGAGATTCGCCTTAGTCTCGTGGGCTCGGAGATGT GTATAAGAGACAGCAAGCGTTGGCTTCTCGCATCT |
| N702 | CAAGCAGAAGACGGCATACGAGATCTAGTACGGTCTCGTGGGCTCGGAGATGT GTATAAGAGACAGCAAGCGTTGGCTTCTCGCATCT |
| N703 | CAAGCAGAAGACGGCATACGAGATTTCTGCCTGTCTCGTGGGCTCGGAGATGT GTATAAGAGACAGCAAGCGTTGGCTTCTCGCATCT |

**Table S3. DNA barcode A sequence used in fungal-DBiT.**

| Sequence Name | Sequence (5'-3') |
|---|---|
| Barcode A-1 | /5Phos/AGGCCAGAGCATTCGAACGTGATGTGGCCGATGTTTCG |
| Barcode A-2 | /5Phos/AGGCCAGAGCATTCGAAACATCGGTGGCCGATGTTTCG |
| Barcode A-3 | /5Phos/AGGCCAGAGCATTCGATGCCTAAGTGGCCGATGTTTCG |
| Barcode A-4 | /5Phos/AGGCCAGAGCATTCGAGTGGTCAGTGGCCGATGTTTCG |
| Barcode A-5 | /5Phos/AGGCCAGAGCATTCGACCACTGTGTGGCCGATGTTTCG |
| Barcode A-6 | /5Phos/AGGCCAGAGCATTCGACATTGGCGTGGCCGATGTTTCG |
| Barcode A-7 | /5Phos/AGGCCAGAGCATTCGCAGATCTGGTGGCCGATGTTTCG |
| Barcode A-8 | /5Phos/AGGCCAGAGCATTCGCATCAAGTGTGGCCGATGTTTCG |
| Barcode A-9 | /5Phos/AGGCCAGAGCATTCGCGCTGATCGTGGCCGATGTTTCG |
| Barcode A-10 | /5Phos/AGGCCAGAGCATTCGACAAGCTAGTGGCCGATGTTTCG |
| Barcode A-11 | /5Phos/AGGCCAGAGCATTCGCTGTAGCCGTGGCCGATGTTTCG |
| Barcode A-12 | /5Phos/AGGCCAGAGCATTCGAGTACAAGGTGGCCGATGTTTCG |

| | |
|---|---|
| Barcode A-13 | /5Phos/AGGCCAGAGCATTCGAACAACCAGTGGCCGATGTTTCG |
| Barcode A-14 | /5Phos/AGGCCAGAGCATTCGAACCGAGAGTGGCCGATGTTTCG |
| Barcode A-15 | /5Phos/AGGCCAGAGCATTCGAACGCTTAGTGGCCGATGTTTCG |
| Barcode A-16 | /5Phos/AGGCCAGAGCATTCGAAGACGGAGTGGCCGATGTTTCG |
| Barcode A-17 | /5Phos/AGGCCAGAGCATTCGAAGGTACAGTGGCCGATGTTTCG |
| Barcode A-18 | /5Phos/AGGCCAGAGCATTCGACACAGAAGTGGCCGATGTTTCG |
| Barcode A-19 | /5Phos/AGGCCAGAGCATTCGACAGCAGAGTGGCCGATGTTTCG |
| Barcode A-20 | /5Phos/AGGCCAGAGCATTCGACCTCCAAGTGGCCGATGTTTCG |
| Barcode A-21 | /5Phos/AGGCCAGAGCATTCGACGCTCGAGTGGCCGATGTTTCG |
| Barcode A-22 | /5Phos/AGGCCAGAGCATTCGACGTATCAGTGGCCGATGTTTCG |
| Barcode A-23 | /5Phos/AGGCCAGAGCATTCGACTATGCAGTGGCCGATGTTTCG |
| Barcode A-24 | /5Phos/AGGCCAGAGCATTCGAGAGTCAAGTGGCCGATGTTTCG |
| Barcode A-25 | /5Phos/AGGCCAGAGCATTCGAGATCGCAGTGGCCGATGTTTCG |
| Barcode A-26 | /5Phos/AGGCCAGAGCATTCGAGCAGGAAGTGGCCGATGTTTCG |
| Barcode A-27 | /5Phos/AGGCCAGAGCATTCGAGTCACTAGTGGCCGATGTTTCG |
| Barcode A-28 | /5Phos/AGGCCAGAGCATTCGATCCTGTAGTGGCCGATGTTTCG |
| Barcode A-29 | /5Phos/AGGCCAGAGCATTCGATTGAGGAGTGGCCGATGTTTCG |
| Barcode A-30 | /5Phos/AGGCCAGAGCATTCGCAACCACAGTGGCCGATGTTTCG |
| Barcode A-31 | /5Phos/AGGCCAGAGCATTCGGACTAGTAGTGGCCGATGTTTCG |
| Barcode A-32 | /5Phos/AGGCCAGAGCATTCGCAATGGAAGTGGCCGATGTTTCG |
| Barcode A-33 | /5Phos/AGGCCAGAGCATTCGCACTTCGAGTGGCCGATGTTTCG |
| Barcode A-34 | /5Phos/AGGCCAGAGCATTCGCAGCGTTAGTGGCCGATGTTTCG |
| Barcode A-35 | /5Phos/AGGCCAGAGCATTCGCATACCAAGTGGCCGATGTTTCG |
| Barcode A-36 | /5Phos/AGGCCAGAGCATTCGCCAGTTCAGTGGCCGATGTTTCG |
| Barcode A-37 | /5Phos/AGGCCAGAGCATTCGCCGAAGTAGTGGCCGATGTTTCG |
| Barcode A-38 | /5Phos/AGGCCAGAGCATTCGCCGTGAGAGTGGCCGATGTTTCG |
| Barcode A-39 | /5Phos/AGGCCAGAGCATTCGCCTCCTGAGTGGCCGATGTTTCG |
| Barcode A-40 | /5Phos/AGGCCAGAGCATTCGCGAACTTAGTGGCCGATGTTTCG |
| Barcode A-41 | /5Phos/AGGCCAGAGCATTCGCGACTGGAGTGGCCGATGTTTCG |
| Barcode A-42 | /5Phos/AGGCCAGAGCATTCGCGCATACAGTGGCCGATGTTTCG |
| Barcode A-43 | /5Phos/AGGCCAGAGCATTCGCTCAATGAGTGGCCGATGTTTCG |
| Barcode A-44 | /5Phos/AGGCCAGAGCATTCGCTGAGCCAGTGGCCGATGTTTCG |
| Barcode A-45 | /5Phos/AGGCCAGAGCATTCGCTGGCATAGTGGCCGATGTTTCG |
| Barcode A-46 | /5Phos/AGGCCAGAGCATTCGGAATCTGAGTGGCCGATGTTTCG |
| Barcode A-47 | /5Phos/AGGCCAGAGCATTCGCAAGACTAGTGGCCGATGTTTCG |
| Barcode A-48 | /5Phos/AGGCCAGAGCATTCGGAGCTGAAGTGGCCGATGTTTCG |
| Barcode A-49 | /5Phos/AGGCCAGAGCATTCGGATAGACAGTGGCCGATGTTTCG |
| Barcode A-50 | /5Phos/AGGCCAGAGCATTCGGCCACATAGTGGCCGATGTTTCG |

**Table S4. DNA barcodeB sequence used in fungal-DBiT.**

| Sequence Name | Sequence (5'-3') |
|---|---|
| barcode B-1 | CAAGCGTTGGCTTCTCGCATCTAACGTGATATCCACGTGCTTGAG |
| barcode B-2 | CAAGCGTTGGCTTCTCGCATCTAAACATCGATCCACGTGCTTGAG |
| barcode B-3 | CAAGCGTTGGCTTCTCGCATCTATGCCTAAATCCACGTGCTTGAG |
| barcode B-4 | CAAGCGTTGGCTTCTCGCATCTAGTGGTCAATCCACGTGCTTGAG |
| barcode B-5 | CAAGCGTTGGCTTCTCGCATCTACCACTGTATCCACGTGCTTGAG |
| barcode B-6 | CAAGCGTTGGCTTCTCGCATCTACATTGGCATCCACGTGCTTGAG |

| | |
|---|---|
| barcode B-7 | CAAGCGTTGGCTTCTCGCATCTCAGATCTGATCCACGTGCTTGAG |
| barcode B-8 | CAAGCGTTGGCTTCTCGCATCTCATCAAGTATCCACGTGCTTGAG |
| barcode B-9 | CAAGCGTTGGCTTCTCGCATCTCGCTGATCATCCACGTGCTTGAG |
| barcode B-10 | CAAGCGTTGGCTTCTCGCATCTACAAGCTAATCCACGTGCTTGAG |
| barcode B-11 | CAAGCGTTGGCTTCTCGCATCTCTGTAGCCATCCACGTGCTTGAG |
| barcode B-12 | CAAGCGTTGGCTTCTCGCATCTAGTACAAGATCCACGTGCTTGAG |
| barcode B-13 | CAAGCGTTGGCTTCTCGCATCTAACAACCAATCCACGTGCTTGAG |
| barcode B-14 | CAAGCGTTGGCTTCTCGCATCTAACCGAGAATCCACGTGCTTGAG |
| barcode B-15 | CAAGCGTTGGCTTCTCGCATCTAACGCTTAATCCACGTGCTTGAG |
| barcode B-16 | CAAGCGTTGGCTTCTCGCATCTAAGACGGAATCCACGTGCTTGAG |
| barcode B-17 | CAAGCGTTGGCTTCTCGCATCTAAGGTACAATCCACGTGCTTGAG |
| barcode B-18 | CAAGCGTTGGCTTCTCGCATCTACACAGAAATCCACGTGCTTGAG |
| barcode B-19 | CAAGCGTTGGCTTCTCGCATCTACAGCAGAATCCACGTGCTTGAG |
| barcode B-20 | CAAGCGTTGGCTTCTCGCATCTACCTCCAAATCCACGTGCTTGAG |
| barcode B-21 | CAAGCGTTGGCTTCTCGCATCTACGCTCGAATCCACGTGCTTGAG |
| barcode B-22 | CAAGCGTTGGCTTCTCGCATCTACGTATCAATCCACGTGCTTGAG |
| barcode B-23 | CAAGCGTTGGCTTCTCGCATCTACTATGCAATCCACGTGCTTGAG |
| barcode B-24 | CAAGCGTTGGCTTCTCGCATCTAGAGTCAAATCCACGTGCTTGAG |
| barcode B-25 | CAAGCGTTGGCTTCTCGCATCTAGATCGCAATCCACGTGCTTGAG |
| barcode B-26 | CAAGCGTTGGCTTCTCGCATCTAGCAGGAAATCCACGTGCTTGAG |
| barcode B-27 | CAAGCGTTGGCTTCTCGCATCTAGTCACTAATCCACGTGCTTGAG |
| barcode B-28 | CAAGCGTTGGCTTCTCGCATCTATCCTGTAATCCACGTGCTTGAG |
| barcode B-29 | CAAGCGTTGGCTTCTCGCATCTATTGAGGAATCCACGTGCTTGAG |
| barcode B-30 | CAAGCGTTGGCTTCTCGCATCTCAACCACAATCCACGTGCTTGAG |
| barcode B-31 | CAAGCGTTGGCTTCTCGCATCTGACTAGTAATCCACGTGCTTGAG |
| barcode B-32 | CAAGCGTTGGCTTCTCGCATCTCAATGGAAATCCACGTGCTTGAG |
| barcode B-33 | CAAGCGTTGGCTTCTCGCATCTCACTTCGAATCCACGTGCTTGAG |
| barcode B-34 | CAAGCGTTGGCTTCTCGCATCTCAGCGTTAATCCACGTGCTTGAG |
| barcode B-35 | CAAGCGTTGGCTTCTCGCATCTCATACCAAATCCACGTGCTTGAG |
| barcode B-36 | CAAGCGTTGGCTTCTCGCATCTCCAGTTCAATCCACGTGCTTGAG |
| barcode B-37 | CAAGCGTTGGCTTCTCGCATCTCCGAAGTAATCCACGTGCTTGAG |
| barcode B-38 | CAAGCGTTGGCTTCTCGCATCTCCGTGAGAATCCACGTGCTTGAG |
| barcode B-39 | CAAGCGTTGGCTTCTCGCATCTCCTCCTGAATCCACGTGCTTGAG |
| barcode B-40 | CAAGCGTTGGCTTCTCGCATCTCGAACTTAATCCACGTGCTTGAG |
| barcode B-41 | CAAGCGTTGGCTTCTCGCATCTCGACTGGAATCCACGTGCTTGAG |
| barcode B-42 | CAAGCGTTGGCTTCTCGCATCTCGCATACAATCCACGTGCTTGAG |
| barcode B-43 | CAAGCGTTGGCTTCTCGCATCTCTCAATGAATCCACGTGCTTGAG |
| barcode B-44 | CAAGCGTTGGCTTCTCGCATCTCTGAGCCAATCCACGTGCTTGAG |
| barcode B-45 | CAAGCGTTGGCTTCTCGCATCTCTGGCATAATCCACGTGCTTGAG |
| barcode B-46 | CAAGCGTTGGCTTCTCGCATCTGAATCTGAATCCACGTGCTTGAG |
| barcode B-47 | CAAGCGTTGGCTTCTCGCATCTCAAGACTAATCCACGTGCTTGAG |
| barcode B-48 | CAAGCGTTGGCTTCTCGCATCTGAGCTGAAATCCACGTGCTTGAG |
| barcode B-49 | CAAGCGTTGGCTTCTCGCATCTGATAGACAATCCACGTGCTTGAG |
| barcode B-50 | CAAGCGTTGGCTTCTCGCATCTGCCACATAATCCACGTGCTTGAG |
